# BOLD signal changes can oppose oxygen metabolism across the human cortex

**DOI:** 10.1101/2023.12.08.570806

**Authors:** Samira M. Epp, Gabriel Castrillón, Beijia Yuan, Jessica Andrews-Hanna, Christine Preibisch, Valentin Riedl

## Abstract

Functional MRI measures brain activity by monitoring changes in blood oxygen levels, known as the blood-oxygen-level-dependent (BOLD) signal, rather than measuring neuronal activity directly. This approach crucially relies on neurovascular coupling, the mechanism that links neuronal activity to changes in cerebral blood flow (CBF). However, it remains unclear whether this relationship is consistent for both positive and negative BOLD responses across the human cortex. Here we found that about 40% of voxels with significant BOLD signal changes during various tasks showed reversed oxygen metabolism, particularly in the default mode network. These “discordant” voxels differed in baseline oxygen extraction fraction (OEF) and regulated oxygen demand via OEF changes, while “concordant” voxels depended mainly on CBF changes. Our findings challenge the canonical interpretation of the BOLD signal, indicating that quantitative fMRI provides a more reliable assessment of both absolute and relative changes in neuronal activity.

## INTRODUCTION

Neuronal activity is the primary energy consumer in the brain, driven by oxygen metabolism and quantified as the cerebral metabolic rate of oxygen (CMRO_2_) ^1^. Functional magnetic resonance imaging (fMRI) maps this activity indirectly by detecting regional changes in blood oxygenation ^2^. The resulting blood-oxygenation-level-dependent (BOLD) signal originates from fluctuations in deoxygenated hemoglobin, rather than from neuronal activity itself.

Interpreting BOLD signal changes (ΔBOLD) as changes in neuronal activity depends on neurovascular coupling, the process that links neuronal activity to local changes in cerebral blood flow (CBF) ^3–7^. Classical work in human sensory cortices using positron emission tomography (PET) showed that sensory stimulation evokes modest increases in CMRO_2_ but a disproportionately larger increase in CBF, resulting in a positive coupling ratio of ΔCBF/ΔCMRO_2_ (*n*-ratio ∼ 2 - 4) ^8^. This surplus in CBF forms the basis of the canonical hemodynamic response (*n*-ratio > 1), which generally allows the interpretation of a positive BOLD response as increased neuronal activity, and vice versa ^9,10^. Mesoscopic studies have supported this principle by linking heightened neural or synaptic activity to increased CBF and positive BOLD responses ^11–13^. Likewise, others have shown that inhibitory processes and decreased activity are linked to negative BOLD changes ^14–18^. However, it remains unclear whether the canonical hemodynamic response applies uniformly beyond sensory cortices, particularly throughout the entire human cortex.

Several studies have shown that BOLD signal changes do not always accurately reflect neuronal activity. Animal studies identified task-induced changes in CBF and metabolic activity, accompanied by minimal or opposite BOLD signal responses ^19,20^, indicating conditions where hemodynamics and neuronal activity are decoupled ^4,21^. Reports of unchanged or even increased metabolism despite significant negative ΔBOLD or ΔCBF further challenge the assumption of uniform neurovascular coupling ^22–25^. The BOLD signal itself reflects a complex interplay among changes in CBF, cerebral blood volume (CBV), and the oxygen extraction fraction (OEF) during capillary passage, making its interpretation region-dependent ^10,26^. Consequently, various studies have reported inconsistencies between BOLD signal responses and cognitive or neuronal activity in humans ^25,27–30^. Moreover, variations in vasculature ^31^ and hemodynamic responsiveness ^9,31–34^ can produce similar macroscopic BOLD patterns through distinct underlying mechanisms. These differences particularly affect the interpretation of BOLD signal patterns in patients with altered hemodynamics ^10,30^. Together, these studies challenge the reliability of the BOLD signal response as an indicator of neuronal activity across the cortex, motivating a more quantitative examination of neurovascular coupling.

We addressed this question by measuring absolute oxygen metabolism and individual vascular components underlying positive and negative BOLD signal changes. The gold standard for CBF and CMRO_2_ measurements is ^15^O PET; yet, this technique requires an on-site cyclotron, a sophisticated imaging setup, and significant experience in handling three different radiotracers (e.g., CBF: 15O-water, CBV: 15O-CO, OEF: 15O-gas) of short half-lives ^8,35^. Furthermore, this invasive method poses significant demands on participants owing to the exposure to radioactivity and arterial sampling. As an alternative, various MRI approaches have been developed over the past three decades to quantify voxel-wise oxygen extraction and metabolism in the human cortex ^36–38^. The quantitative BOLD (qBOLD) approach is based on an analytical model ^39^ that relates R2’ to OEF ^40–42^. A multiparametric variant of the qBOLD technique (mqBOLD) combines separate measurements of the intrinsic and effective relaxation times, T2 and T2* for R2’, with an independent quantification of CBV for OEF measurement. mqBOLD imaging has been widely applied to study patient groups with vascular pathologies and brain tumors ^42–47^, summarized recently by Alzaidi et al. ^36^.

In this study, we used both quantitative and conventional BOLD imaging to test the hypothesis that ΔBOLD would not reliably reflect changes in oxygen metabolism throughout the entire cortex. We found that, in a substantial fraction of voxels with significant BOLD responses, oxygen metabolism changes in the opposite direction to both positive and negative BOLD signals. Notably, these discordant voxels regulated oxygen demand primarily via changes in OEF, rather than CBF. These findings challenge the canonical hemodynamic response model, demonstrating that BOLD signal changes alone can lead to misleading interpretations of underlying neuronal activity.

## RESULTS

Brain imaging data were collected using BOLD and quantitative fMRI while participants completed four experimental conditions within a single session (Figure 1A, 1B). These conditions allowed identification of (1) ‘task-positive’ regions with increased BOLD signal and (2) ‘task-negative’ regions with negative BOLD response during calculation (CALC). We also examined (3) positive BOLD responses in the ‘task-negative’ regions during an autobiographical memory task (MEM). BOLD-contrast images for CALC and MEM were calculated relative to a control task (CTRL) or resting state (REST).

**Figure 1.**
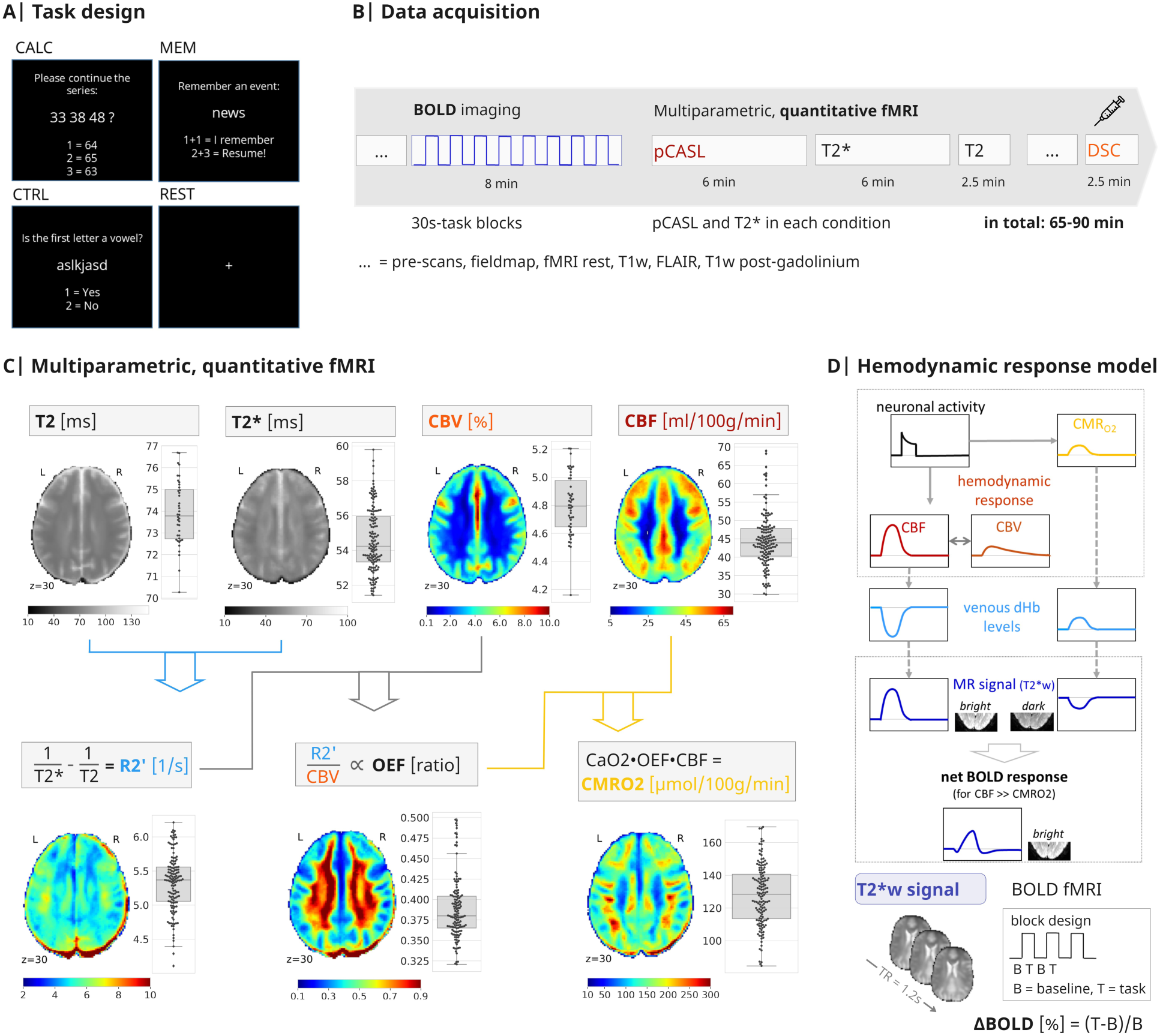
Study design, quantitative fMRI, and the hemodynamic response model of BOLD fMRI. A| We performed BOLD- and quantitative fMRI on healthy subjects performing four different conditions within the same imaging session: calculation task (CALC), autobiographical memory task (MEM), active control baseline (CTRL), and uncontrolled resting-state baseline (REST). **B|** During each imaging session, we acquired BOLD fMRI data with a 30-sec block design, together with multiparametric, quantitative fMRI data while conditions were presented in pseudo-randomized order: T2* mapping to measure both reversible and irreversible dephasing; pseudo-continuous arterial spin labeling (pCASL) to measure CBF during each condition; T2 mapping to once measure irreversible dephasing due to tissue properties; dynamic susceptibility contrast (DSC) MRI using a contrast agent to measure CBV during CTRL, at the end of the session. **C|** Quantitative fMRI delivers voxelwise CMRO_2_ by integrating T2, T2*, CBV, and CBF parameters via Fick’s principle (see Methods section for detailed equations). Brain slices of subject-average parameter maps and boxplots illustrating average GM values across all 40 subjects and all conditions of the main study (line, median; box limits, upper and lower quartiles; whiskers, minimum and maximum data points except for outliers: values outside of 1.5* IQR; individual dots, subject average per condition). The reversible transverse relaxation rate (R2’) reflects a voxel’s overall deoxyhemoglobin (dHb) content. The OEF is proportional to R2’/CBV. Voxel-wise CMRO_2_ is then calculated as the product of OEF, CBF, and the arterial oxygen content of the blood (CaO2) as derived from individual measures of oxygen saturation and hematocrit. **D|** The canonical hemodynamic response model of the BOLD signal: Increased neuronal activity leads to higher oxygen metabolism (CMRO_2_), which results in decreased oxygen levels and an increased concentration of dHb in venous blood. Neurovascular coupling mechanisms initiate an increase in CBV and CBF. This hemodynamic response counteracts the effects of oxygen metabolism, ultimately leading to a decrease in dHb content. In BOLD-fMRI, the net fluctuation of dHb levels is dynamically measured with T2*-weighted (T2*w) echo planar imaging (EPI). ΔBOLD during any task condition is calculated by subtracting task (T) from baseline (B) T2*w data. Based on the canonical hemodynamic response function, positive BOLD signal changes are commonly interpreted as increased neuronal activity.

Quantitative fMRI, combining mqBOLD and pCASL MRI, estimates CMRO₂ as a metabolic marker of neuronal activity. CMRO₂ was calculated using Fick’s principle, integrating parameter maps of several aspects of the hemodynamic response into a measure of oxygen metabolism (Figure 1C, Methods). Table 1 lists parameter and CMRO₂ values averaged across all subjects and conditions.

**Table 1.** Whole-brain (GM) parameter values, mean ± SD across 40 subjects and all conditions per subject.

| R2' | CBV | OEF | CBF | CMRO<sub>2</sub> |
| --- | --- | --- | --- | --- |
| [1/s] | [%vol] | [ratio] | [ml/100g/min] | [μmol/100g/min] |
| 5.3 ± 0.4 | 4.8 ± 0.2 | 0.39 ± 0.04 | 44.5 ± 7.2 | 127.8 ± 18.4 |

The canonical hemodynamic response model (Figure 1D) assumes an identical hemodynamic response across voxels, with ΔCBF exceeding ΔCMRO₂ (n-ratio >1). In this study, we used quantitative fMRI to measure hemodynamic parameters per condition and compared them to task-induced BOLD changes relative to an experimental baseline (ΔBOLD [%]).

### Average negative BOLD signal response does not indicate reduced oxygen metabolism

We performed separate partial least squares (PLS) analyses on BOLD and mqBOLD parameter maps to compare BOLD and quantitative fMRI results. Bootstrap ratios were used for statistical mapping (Fig. 2A shows CALC vs. CTRL; additional contrasts in Fig. S2). We found widespread significant positive and negative ΔBOLD for CALC vs. CTRL. Group histograms (Fig. 2B) show task-related changes for each parameter within significant ΔBOLD regions (CALC-positive, CALC-negative). In CALC-positive masks, ΔCBF and ΔCMRO₂ showed canonical amplitudes: ΔBOLD[%] = 0.37%, ΔCBF[%] = 6.5%, ΔCMRO_2_ [%] = 3.1%, *n*-ratio (ΔCBF[%] / ΔCMRO_2_ [%]) = 2.1 (Fig. 2B). CALC-negative masks showed near-zero ΔCBF and ΔCMRO₂, despite negative ΔBOLD reaching 70% of the positive amplitude (−0.26%). To address normalization distortions, we also analyzed native-space data using individual BOLD masks, confirming robust positive ΔCBF and ΔCMRO₂ in CALC-positive masks, with no significant response in CALC-negative masks across subjects (Fig. 2C). A second-level GLM model (Fig. S1A and S1B), instead of the PLS approach confirmed near zero changes of median ΔCBF and ΔCMRO₂ in CALC-negative masks. Overall, positive ΔBOLD values matched the canonical hemodynamic response, with ΔCBF (in %) being at least ten times higher than ΔBOLD and twice the amplitude of ΔCMRO₂ (n-ratio >2). However, no significant negative ΔCBF, ΔCBV, ΔT2*, or ΔCMRO₂ were found in negative ΔBOLD masks across subjects.

**Figure 2.**
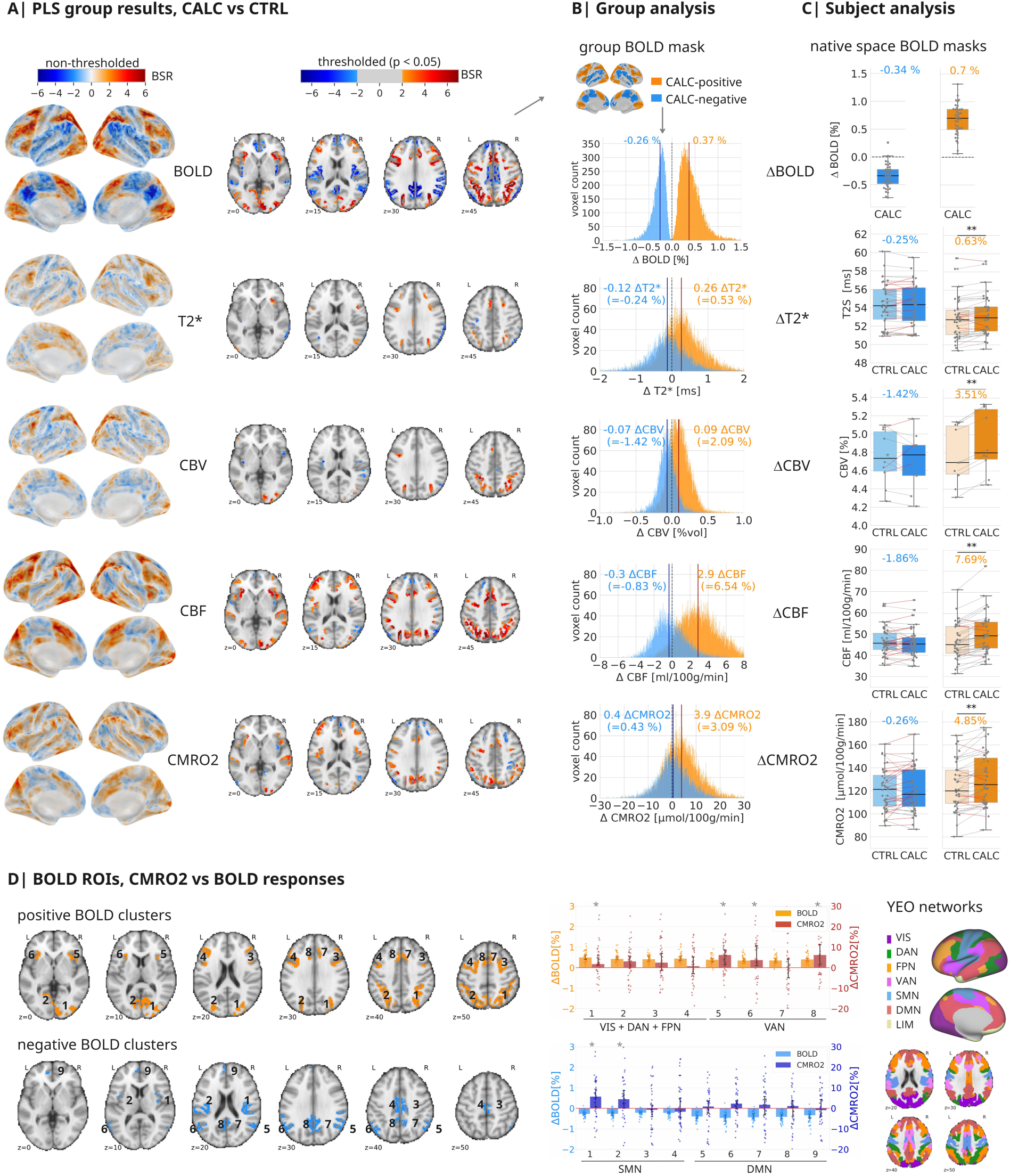
Negative BOLD signal changes do not indicate reduced oxygen metabolism. All analyses across N=40 subjects of the main study, CBV results across N=10. A| Statistical results of the PLS group analysis in standard MNI space, comparing CALC and CTRL, for both BOLD and quantitative fMRI parameter maps. Maps projected on the brain surface show non-thresholded bootstrap ratios (BSR, akin to z-values) of the first latent variable, which was significant in each analysis, i.e., for BOLD, CBF, CBV (N=10), T2* and CMRO_2_ respectively (permutation tests, *p* < .001). Red voxels indicate higher values during CALC compared to CTRL, while blue voxels represent the opposite contrast. Axial slices display significant brain clusters, thresholded at BSR ±2 (akin to *p* < .05, cluster size > 30 voxels) for each parameter. **B|** Histograms depict group-average median voxel distributions of all parameters (CALC minus CTRL) within the binarized CALC-positive and CALC-negative group masks (shown as surface plot). **C|** Subject distribution of all parameters in native space. Dots represent individual subject values, calculated as the median across voxels within individual BOLD masks of significant positive (orange) and negative (blue) ΔBOLD. Boxplots: line, median; box limits, upper and lower quartiles; whiskers, minimum and maximum data points except for outliers: values outside of 1.5* IQR; individual dots, one dot per subject, median voxel values. Paired samples, two-sided t-tests, ** *p* < .001. Gray lines connect values obtained from individual subjects, while red lines indicate subjects where the direction of ΔCBF or ΔCMRO_2_ deviates from what the BOLD-signal suggests. **D|** Regional clusters (designated by numbers) of positive and negative ΔBOLD (PLS group results, thresholded at BSR > ±3) and associated bar plots showing ΔBOLD and ΔCMRO_2_. Please note the different axes for ΔBOLD and ΔCMRO_2_. Bars indicate ΔBOLD[%] and ΔCMRO_2_[%] median across voxels, error bars 95% CI, 2000 bootstraps, and dots represent subject values. Any signal increase or decrease for which error bars cross the zero line is considered non-significant; an asterisk denotes *p* < .05, not corrected for multiple comparisons. Clusters are located in different functional networks, as depicted on the brain surface and in axial slices ^79^: visual (VIS); dorsal attention (DAN); frontoparietal (FPN); ventral attention (VAN); somatomotor (SMN); default mode (DMN) network.

We then analyzed metabolic parameters regionally, avoiding brain-wide averaging effects. Eight clusters showed positive, nine clusters negative ΔBOLD (Fig. 2D, left). Median ΔCMRO₂ was significantly positive in half of the positive ΔBOLD clusters, but negative in none of the negative clusters (Fig. 2D, bar plots). Notably, two auditory network clusters (SMN 1, SMN 2) with negative ΔBOLD had significant positive ΔCMRO₂.

### Control analyses

We also examined ΔBOLD and ΔCMRO₂ for the other contrasts, CALC vs. REST (Fig. S2A-C) and MEM vs. CTRL (Fig. S2D-G). CALC-positive and CALC-negative masks showed higher overall ΔBOLD, ΔT2*, ΔCBF, and ΔCMRO₂ compared to CTRL, but changes within CALC-negative masks remained nonsignificant (Fig. S2C). For the MEM task, results mirrored the CALC patterns, with similar BOLD signal amplitudes, and stronger positive than negative ΔBOLD. However, ΔCBF and ΔCMRO₂ were only significant in the MEM-positive mask. Thus, positive ΔBOLD reliably indicated significant positive ΔCMRO₂ across conditions, but the metabolic interpretation of negative ΔBOLD remained unclear. Table S1 summarizes average parameter changes for all conditions.

Several factors could explain the absence of hemodynamic and metabolic changes, especially related to negative ΔBOLD: (i) voxel-specific signal confounds, (ii) limited sensitivity for detecting negative ΔCBF or ΔCMRO₂, or (iii) distinct hemodynamic mechanisms in certain regions. We address each explanation below.

Most MEM-positive voxels overlapped with the default mode network (DMN), which showed negative ΔBOLD during CALC (Fig. S2F). To test for voxel-specific confounds, we analyzed CBF and CMRO₂ in voxels occurring in both MEM-positive and CALC-negative masks. Fig. S2H shows the histograms of ΔBOLD, ΔCBF, and ΔCMRO_2_ for all voxels of a conjunction mask (MEM-positive ∧ CALC-negative). For positive ΔBOLD (0.32%), we observed a canonical increase in ΔCBF (5.94%) and ΔCMRO₂ (3.54%). However, in the same voxels, a negative ΔBOLD (−0.29%) was associated with a much weaker ΔCBF (−0.47%), resulting in a positive ΔCMRO₂ (1.68%). Thus, a lower ΔCBF than expected from the canonical framework results in elevated ΔCMRO₂ for negative ΔBOLD. Crucially, the absence of canonical negative BOLD responses in “conjunction voxels” cannot be explained by voxel-specific confounds, as those same voxels do display a canonical response for positive ΔBOLD.

According to the hemodynamic response model (Fig. 1D), neurovascular coupling is primarily driven by changes in CBF. To validate the sensitivity of pCASL imaging for detecting ΔCBF, we performed several control analyses (Fig. S3). We found ΔCBF magnitudes ranging from −12% to +29%, with regions of positive ΔCBF showing higher n-ratios than those with negative ΔCBF (Fig. S3A). Moreover, the spatial pattern of negative ΔCBF during CALC matched that observed with PET from a different study (Fig. S3B). Notably, CBF decreases identified by PET and MRI were more localized than BOLD decreases and centered on peak regions. Additionally, we also validated the homogeneity of our MR-derived OEF with independent PET data (Fig. S3C).

We also assessed signal stability of BOLD and CBF measurements during continuous task performance. In the main study (Fig. 1), block durations vary for BOLD (30 sec) and quantitative (approx. 6 min) fMRI to achieve the best signal-to-noise ratio (SNR) for each modality. In a control study (N=18), participants performed 3-minute CALC and MEM blocks while BOLD data were collected. BOLD signals remained stable without habituation or drift (Fig. S3D), with amplitude changes comparable to the main study (CALC-positive/negative/MEM-positive: 0.6%/-0.46%/0.34% vs. 0.70%/-0.34%/0.51%). Time-resolved pCASL analysis from the main study also showed constant CBF data throughout measurement blocks (Fig. S3E).

### High prevalence of discordant voxels among positive and negative BOLD responses

We next assessed the ΔBOLD–ΔCMRO₂ relationships on the voxel level and compared our data to an established model of hemodynamic responses. The Davis model predicts ΔBOLD responses based on realistic cortical ranges for ΔCMRO_2_[%] and ΔCBF[%] (Davis et al., 1998). Fig. 3A displays the model’s prediction of ΔBOLD (contours, −3% to +3%) for a set of typical hemodynamic response parameters. A canonical BOLD response occurs when ΔCBF exceeds ΔCMRO₂ (n-ratio >1). However, the model also predicts discordant responses – where ΔBOLD disagrees with metabolic change – when ΔCBF is lower than ΔCMRO₂ (n-ratio <1) or their signs differ (Fig. 3A, violet shadings). For example, a negative ΔBOLD of −1.3% would arise if CBF increases by only about 5% during a ΔCMRO_2_ of +23%. In other words, the same metabolic change can result in opposite signs of the BOLD signal, depending on the magnitude of related ΔCBF. Since we measured all relevant parameters and calculated ΔCMRO₂ using Fick’s principle, we next compared our empirical data to the model predictions.

**Figure 3.**
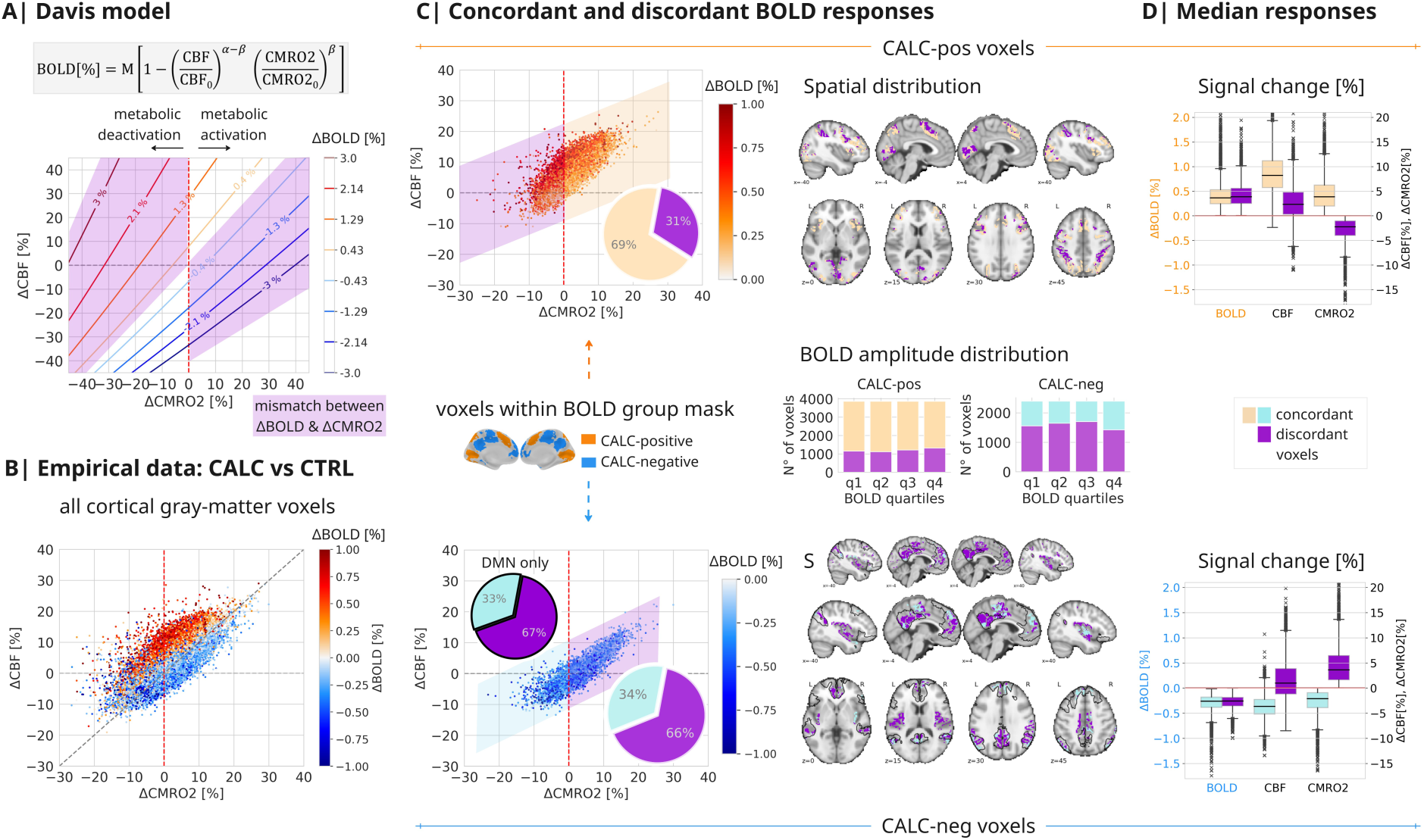
Dependence of BOLD signal responses on changes in CBF and CMRO_2_. A| Visualization of the Davis model. Predicted BOLD responses (ΔBOLD[%]) are depicted as colored contour lines across ΔCBF[%] and ΔCMRO_2_[%]. The model illustrates that for a given ΔCMRO_2_[%], the direction and magnitude of the BOLD response depend on the amplitudes of ΔCBF[%]. The canonical hemodynamic response assumes that an increase in ΔCBF will exceed the increase in ΔCMRO_2_, resulting in a positive ΔBOLD, and vice versa for negative BOLD signal changes. However, the Davis model also predicts a range of BOLD responses with discordant signs compared to the changes in underlying metabolic activity (violet shading). **B|** Empirical data from BOLD- and quantitative fMRI for all cortical voxels, median across N=40 subjects of the main study. ΔCMRO_2_ was calculated from BOLD-derived ΔR2’ (see Methods), median across all subjects per voxel, with colors indicating ΔBOLD[%]. Our experimental data replicate the slope and range of values predicted by the Davis model, particularly demonstrating a substantial number of voxels with ΔBOLD opposing ΔCMRO_2_. **C|** Voxels, representing the median across N=40 subjects from the main study, display significantly positive (top) and negative (bottom) BOLD responses from (B). Voxels with ΔBOLD concordant to ΔCMRO_2_ are highlighted on a light yellow/turquoise background, while voxels with discordant ΔBOLD values are shown on a violet background. The pie charts illustrate the ratio of discordant to concordant voxels. In the lower panel, the pie chart with black contours specifically summarizes the ratio of discordant voxels within the DMN. Axial slices illustrate the spatial distribution of significantly concordant and discordant BOLD voxels, with black contours indicating the DMN. The stacked bars in the right center illustrate the amplitude distribution of discordant and concordant voxels across BOLD amplitude quartiles. It is noteworthy that discordant voxels show neither spatial nor amplitude preference. **D|** The boxplots summarize ΔCMRO_2_[%] and ΔCBF[%] across all voxels, median across N=40 subjects of the main study, with either concordant or discordant ΔBOLD[%]; (line, median; box limits, upper and lower quartiles; whiskers, minimum and maximum data points except for outliers: values outside of 1.5* IQR; based on median voxel values across subjects). Please note that discordant and concordant voxels exhibit similar ΔBOLD amplitude distributions, even though they signal opposite metabolic responses.

Fig. 3B presents subject-averaged voxel data for CALC vs. CTRL, colored by positive/negative ΔBOLD (Fig. S4A for CALC vs. REST, and Fig. S4B for MEM vs. CTRL). To mitigate low SNR at the voxel level, ΔCMRO₂ was calculated using more stable ΔR2’ data from BOLD-fMRI (see Methods and Fig. S5 for sensitivity analysis). Our empirical data support several predictions of the Davis model: positive ΔBOLD voxels cluster above the line with a slope of 1, indicating an *n*-ratio >1. We also identify a significant number of discordant voxels – those with opposite ΔBOLD and ΔCMRO₂. Those can be summarized as red voxels on the left and blue voxels on the right side of zero Δ CMRO_2_ (Fig. 3B, red dashed line). Interpreting the activity in discordant voxels using only BOLD data and assuming a uniform canonical response would lead to substantial misinterpretation.

To highlight the amount of discordant responses, we plotted CALC-positive (Fig. 3C top) and CALC-negative (Fig. 3C bottom) voxels separately, highlighting discordant voxels (violet shading). Discordant voxels account for 31% and 66% of positive and negative ΔBOLD, respectively (Fig. 3C, pie charts); with a similar ratio of discordant negative ΔBOLD within the DMN (pie chart “DMN only”), the network with the most consistently reported negative BOLD signal responses. Moreover, discordant voxels are spatially distributed across the cortex (Fig. 3C, brain slices) and occur equally among voxels from the lowest to highest quartile of BOLD signal amplitudes (Fig. 3C, bar plots). We repeated these analyses for CALC vs. REST (Fig. S4A) and MEM vs. CTRL (Fig. S4B). Again, discordant voxels appeared with a uniform cortical distribution and across the full range of ΔBOLD, amounting to 36% / 52% of significant positive / negative ΔBOLD voxels in CALC vs. REST and 12% / 54% in MEM vs. CTRL.

Fig. 3D summarizes voxel-median parameter values separately for concordant and discordant voxels. Concordant voxels with either positive or negative ΔBOLD demonstrate a canonical hemodynamic response (*n*-ratio 2.0 for positive, 1.6 for negative ΔBOLD; Fig. 3D, yellow and turquoise bars). Discordant voxels, however, show ΔBOLD opposite to their metabolic response and lower than expected ΔCBF (Fig. 3D, violet bars). Notably, discordant ΔBOLD occurred in about one-third of positive and two-thirds of negative ΔBOLD voxels across different tasks and all magnitude ranges, distributed evenly throughout the cortex.

Potentially, discordance between ΔBOLD and ΔCMRO_2_ could result from partial volume effects when integrating voxel data from imaging sequences with heterogeneous voxel sizes, as occurred in our main study.. To address this, we conducted a replication study (N = 10) using harmonized acquisition matrices for BOLD and mqBOLD data. Results mirrored the main study (Fig. S6): 40% of all voxels with either significant positive or negative ΔBOLD were classified as discordant, and discordant voxels again showed smaller CBF responses than expected from canonical neurovascular coupling.

Finally, we directly estimated the parameters of the Davis model using our own data. With measurements of ΔBOLD, CBV, OEF, and CBF, we calculated α and each subject’s calibration factor M empirically. The average values were α = 0.38 and M = 11.2 ± 1% (N = 40). Fig. S7 replicates Fig. 3 using the Davis model parameters instead of Fick’s formula to quantify CMRO₂.

### ‘Mixed’ voxels switch between concordant and discordant BOLD responses across tasks

Next, we examined whether voxels with both positive and negative BOLD responses during CALC or MEM (using the conjunction mask of the control analysis in Fig. S2H) exhibit consistent hemodynamic response patterns for both signs of ΔBOLD. Voxels were classified as ‘concordant only’, ‘discordant only’, or ‘mixed’ based on whether BOLD and metabolic responses matched in both tasks, differed in both tasks, or showed a mixed pattern (Fig. 4A). The pie chart illustrates that most voxels (48%) were ‘mixed’, while 35% and 18% were consistently concordant or discordant, respectively. Fig. 4B displays the spatial and amplitude distributions for ΔCBF and ΔCMRO₂ across the three categories. ‘Concordant only’ voxels showed canonical hemodynamic responses (*n*-ratio ∼2) regardless of BOLD sign or task (yellow/turquoise bars), while ‘discordant only’ voxels lacked canonical responses for either direction (violet/lilac bars). ‘Mixed’ voxels exhibited canonical coupling for positive BOLD, but not for negative BOLD (red/orange bars). Because changes in CMRO₂, which are not modulated by CBF, must be matched by OEF changes, we tested whether baseline OEF varies by voxel type. We hypothesized that ‘discordant only’ voxels, with the lowest ΔCBF during a task, would have the lowest baseline OEF (indicating the highest oxygen buffer). In line with this hypothesis, baseline OEF was lowest in ‘discordant only’ voxels, intermediate in ‘mixed’, and highest in ‘concordant only’ voxels (Fig. 4C), suggesting that discordant voxels compensate higher oxygen demand mainly via OEF changes. Importantly, OEF data were independently acquired from CBF data.

**Figure 4.**
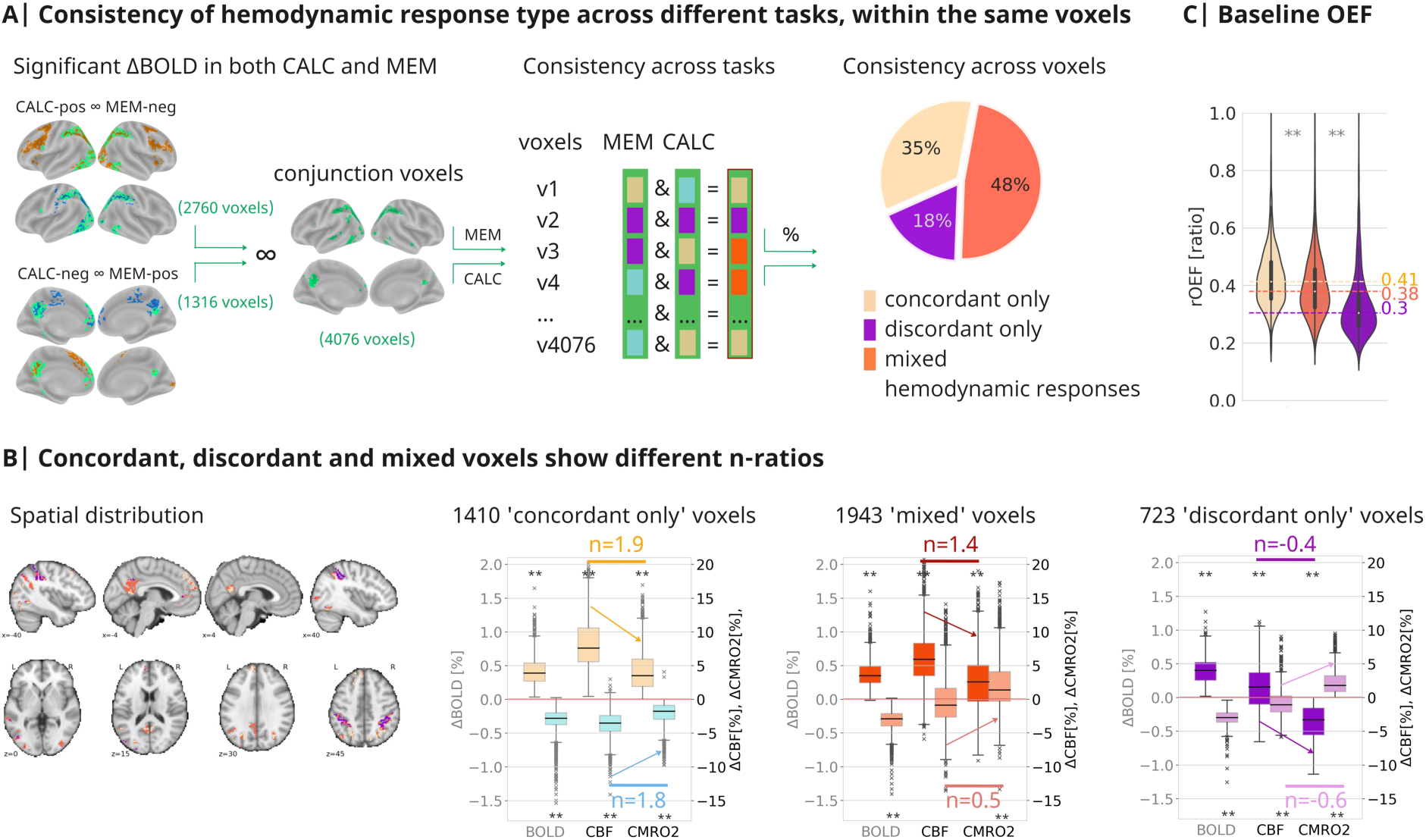
Concordant or discordant BOLD signal responses depend on baseline OEF. Median voxel values across N=30 subjects of the main study. A| Conjunction voxels (green outlines) display both positive and negative changes in BOLD signal during the CALC and MEM conditions. These voxels were categorized based on their response types. The pie chart summarizes the consistency of these response types across voxels: 35% of all conjunction voxels showed concordant responses, meaning they had the same sign of ΔBOLD and ΔCMRO_2_ during both MEM and CALC conditions (‘concordant only’). 18% of all conjunction voxels showed discordant responses in both conditions (‘discordant only’). 48% of the conjunction voxels (‘mixed’ voxels) showed concordant responses in one condition and discordant responses in the other. **B|** The spatial distribution of voxel categories is illustrated on brain slices with consistent color overlays: ‘Concordant only’ and ‘mixed’ voxels primarily cluster in regions of the DMN, while ‘discordant only’ voxels occur in the medial part of the VAN. The boxplots display subject-median parameter values separately for ‘concordant only,’ ‘mixed,’ and ‘discordant only’ voxels (line, median; whiskers, minimum and maximum data points except for outliers: values outside of 1.5* IQR; based on median voxel values across subjects). While the range of ΔBOLD is similar across the three categories, ΔCBF and ΔCMRO_2_ vary strongly as expressed by their *n*-ratios. Asterisks indicate significant differences from zero, tested via independent t-tests, two-sided, corrected for multiple comparisons. **C|** The three voxel categories exhibit significantly different OEF during CTRL baseline, with ‘concordant only’ voxels having significantly higher OEF and ‘discordant only’ voxels having significantly lower OEF than ‘mixed’ voxels. ** *p* < 0.001, independent sample permutation test on the median values, two-sided, conducted for ‘mixed’ versus the two other voxel types, 2000 permutations.

### Baseline OEF predicts different hemodynamic response types

We next examined baseline hemodynamics across the cortex by pooling all voxels with significant ΔBOLD during either task (CALC, MEM) – comprising 53% of all GM voxels – and categorizing them as concordant, discordant, or not task-involved (Fig. 5A, pie chart). Regression plots show that baseline CMRO₂ correlates linearly with CBF, OEF, and CBV across all voxel groups, consistent with Fick’s principle. Multiple linear regression analyses using all three parameters showed that baseline OEF accounted for >68% of CMRO₂ variance, followed by CBF (>28%) and CBV (>1%), with similar model parameters for concordant and discordant voxels (concordant model: F(3,21809) = 72070, *p* < .001, CMRO_2_= 139.9 + 321.1*OEF + 3.3*CBF + 0.4*CBV + e, R^2^=0.91; discordant model: F(3,14544) = 48390, *p* < .001, CMRO_2_= 128.1 + 309.2*OEF + 3.2*CBF + 1.3*CBV + e; R^2^=0.91; all predictors mean-centered, *p*<.001 for all beta values). According to the CMRO_2_ model parameter, discordant voxels have a lower baseline metabolism, while concordant voxels have significantly higher baseline CBF and OEF but lower CBV (Fig. 5B).

**Figure 5.**
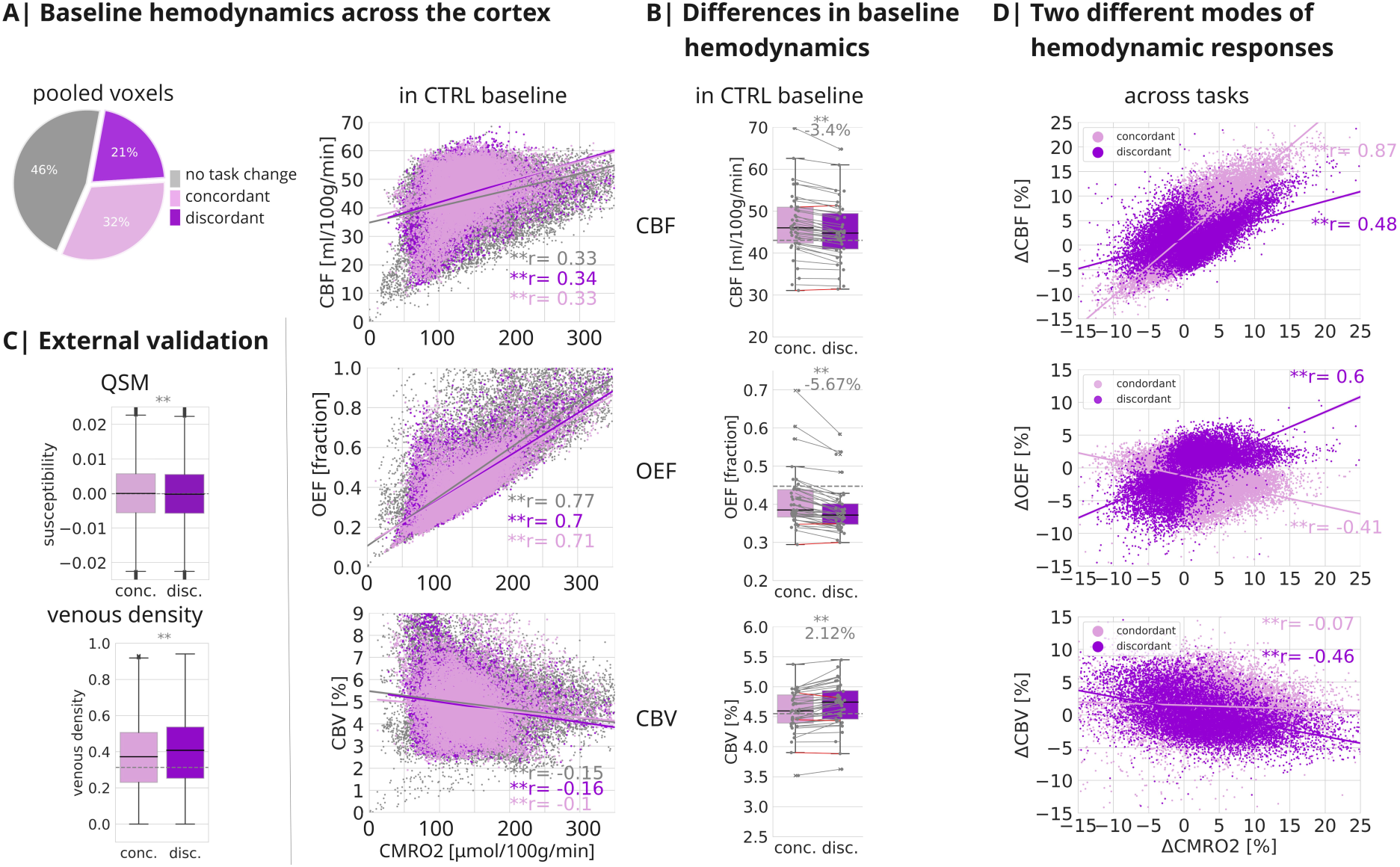
Two types of hemodynamic responses via changes in CBF or OEF. A| Across all task conditions, 21% of all GM voxels showed discordant, 32% showed concordant responses, and 46% were not significantly involved in any task. Regression plots: During baseline, CMRO2 and CBF show a moderate correlation (Pearson’s *r* > 0.32, *p* < .001), CMRO_2_ and OEF are highly correlated (*r* > 0.7, *p* < .001), and CMRO_2_ and CBV show a weak negative correlation (*r* < −0.1, *p* < 0.001) across all groups of voxels, voxel medians across 40 subjects. **B|** Across all 40 subjects, discordant (disc.) voxels, on average, show significantly lower CBF (−3.4%), OEF (−5.7%), and higher CBV (2.1%) than concordant (conc.) voxels (paired-samples, two-sided t-test, *p* < .001). Boxplots: line, median; box limits, upper and lower quartiles; whiskers, minimum and maximum data points except for outliers: values outside of 1.5* IQR; individual dots, median subject values across voxels; gray line, median of remaining GM voxels. **C|** Marginally, but significantly lower baseline susceptibility (QSM) and higher venous density in discordant vs. concordant voxels, revealed by two external datasets ^68,69^. Boxplots: line, median; box limits, upper and lower quartiles; whiskers, minimum and maximum data points except for outliers: values outside of 1.5* IQR; based on voxel values; gray line, median of remaining GM voxels. ** *p* < .001, independent samples permutation test on the median values, two-sided, conducted for discordant versus concordant voxels, 2000 permutations. **D|** Correlation of ΔCMRO_2_[%] with either ΔCBF[%], ΔOEF[%], or ΔCBV[%] across all task contrasts (CALC vs. CTRL: voxel median across 40 subjects, MEM vs. CTRL: voxel median across 30 subjects), separately for concordant (pale) and discordant (dark violet) voxels. ΔCMRO_2_ and ΔCBF showed a strong positive correlation across concordant voxels (*r* = .87, *p* < 0.001) and a moderate correlation across discordant voxels (*r* = .48, *p* < .001). In contrast, ΔCMRO_2_ and ΔOEF showed a moderate, positive correlation across discordant voxels (*r* = .60, *p* < .001) and a weak, negative correlation across concordant voxels (*r* = −.41, *p* < .001). ΔCBV could only be evaluated for a subset of 10 subjects and only across the CALC vs. CTRL contrast, see Methods. ΔCMRO_2_ and ΔCBV showed a weak correlation around zero across concordant voxels (*r* = − .07) and a moderate, negative correlation across discordant voxels (*r* = −.46, *p* < 0.001).

To validate lower OEF and higher CBV in discordant voxels, we analyzed two external QSM datasets (Fig. 5C). Projecting our voxel masks onto these data, we observed that discordant voxels had slightly but significantly lower susceptibility (indicating lower dHB) and significantly higher venous density, supporting our findings.

Finally, we investigated whether ΔOEF might serve as an alternative mechanism to ΔCBF in accommodating oxygen demand during task processing. Regression plots show the linear correlation between ΔCMRO₂ and ΔCBF, ΔOEF, and ΔCBV, separately for concordant and discordant voxels, illustrating the significant contribution of each hemodynamic parameter to oxygen metabolism (Fig. 5D). However, we also identified two distinct patterns of hemodynamic responses. Multiple linear regression analyses revealed that in concordant voxels, ΔCMRO₂ was mainly driven by ΔCBF (concordant model: F(2,21971) = 67980, *p* < .001, with ΔCBF explaining 87% and ΔOEF 16% of a total explained variance of R^2^=0.86 of the model: ΔCMRO_2_[%] = 2.6 + 0.83*ΔOEF[%] + 0.86*ΔCBF[%] + e; all predictors mean-centered, *p*<.001 for all beta values). In discordant voxels ΔOEF was the primary regulator, with ΔCBF secondary (discordant model: F(2,14694) = 39320, *p* < .001, with ΔOEF explaining 58% and ΔCBF 42% of a total explained variance of R^2^=0.84 of the model: ΔCMRO_2_[%] = 1.2 + 1.1*ΔOEF[%] + 0.88*ΔCBF[%] + e; all predictors mean-centered, *p*<.001 for all beta values). The negative ΔCMRO₂ − ΔOEF correlation for concordant voxels reflects compensatory oxygenation during ΔCBF surplus (Fig. 5D, middle plot). CBV was not included in these regressions, as ΔCBV was only acquired for the CALC-CTRL contrast. In summary, concordant voxels regulate CMRO₂ mostly via ΔCBF (87%), whereas discordant voxels rely more on ΔOEF (58%) compared to ΔCBF (42%).

## DISCUSSION

In this study, we evaluated the consistency between changes in the BOLD signal and oxygen metabolism in gray-matter voxels across the human cortex. This finding would support the common interpretation of positive and negative ΔBOLD as indicators of increased or decreased neuronal activity. Contrary to the canonical BOLD response model, approximately 40% of brain voxels with significant BOLD changes exhibited opposing changes in oxygen metabolism. Specifically, voxels with positive ΔBOLD showed decreased ΔCMRO_2_, while those with negative ΔBOLD exhibited increased ΔCMRO_2_. By measuring the BOLD signal, CBF, OEF, and CMRO_2_ in the same session, we uncovered distinct neurovascular mechanisms in regions with concordant versus discordant responses. Discordant voxels primarily regulate oxygen demand via ΔOEF, whereas only concordant voxels display an increase in ΔCBF, aligning with canonical predictions. Moreover, discordant voxels demonstrated lower baseline CMRO_2_ and OEF, indicating that their baseline oxygen supply is sufficient to meet metabolic demands. In conclusion, we identified two distinct hemodynamic responses to neuronal activity changes, influenced by baseline OEF and metabolism.

### Canonical hemodynamic response for average positive ΔBOLD

We combined BOLD with quantitative fMRI to investigate hemodynamic and metabolic changes in relation to BOLD signal changes. Participants performed cognitive tasks eliciting both positive and negative BOLD signal responses across the cortex. Specifically, we aimed to elicit both response types in identical voxels, facilitating a direct comparison of their hemodynamic responses. We employed a cognitively demanding calculation task that induced positive ΔBOLD in various attention-related regions ^49,50^ while concurrently inducing negative ΔBOLD in DMN regions ^50,51^. Additionally, participants undertook an autobiographical memory task known to induce positive ΔBOLD in DMN regions, partially overlapping with those of negative ΔBOLD during CALC ^52,53^. This allowed us to examine opposing ΔBOLD within the same voxels.

Our study also addressed the implications of differing baseline states. While BOLD signal responses to cognitive tasks are usually compared against low-level control tasks ^53–55^, negative ΔBOLD was often reported in comparison to an uncontrolled resting state ^51,56^. Consequently, we included both baseline types into the study design, and all experimental conditions successfully elicited anticipated BOLD signal responses. On average, negative BOLD signal responses constituted approximately 50-70% of the amplitude of positive ΔBOLD (Fig. 2). Additionally, ΔBOLD was larger when compared to REST than to CTRL baseline (Table 2, Fig. S2A-C).

Compared to BOLD fMRI, which requires subtraction analyses between conditions, the mqBOLD approach provides quantitative measures for each condition. During CTRL baseline, we observed average GM values of CMRO_2_ and hemodynamic parameters (see Table 1) consistent with existing literature on the healthy human brain ^35,38,42,51,57–59^. Likewise, task-related changes (see Table S1) aligned with prior findings ^58,60,61^, where ΔCMRO_2_ and ΔCBF exceeded positive ΔBOLD by factors of 10-20, yielding an *n*-ratio of approximately 2 (Fig. 2) and thereby conforming to the canonical hemodynamic response ^10,62^.

### Discordant hemodynamics for negative ΔBOLD and in regions with positive ΔBOLD

Despite a canonical hemodynamic response for mean positive ΔBOLD, we did not find significant hemodynamic or metabolic changes for negative ΔBOLD (see CALC vs CTRL, Fig. 2C) for either task condition (see MEM vs CTRL, Fig. S2G) or when comparing against REST baseline (see Fig. S2C). In a region-specific analysis, we also observed significant deviations from the canonical response in four out of eight clusters with positive ΔBOLD (Fig. 2D). One could argue that voxels with discordant ΔBOLD might suffer from voxel-specific artifacts such as partial volume effects or differences in vasculature, leading to insignificant ΔCBF. To address this issue, we designed our study to achieve significant ΔBOLD during both CALC and MEM tasks in identical voxels, facilitating within-voxel comparisons of ΔBOLD and quantitative measures. Among these voxels (Fig. S2H, ‘conjunction voxels’), we observed only weak ΔCBF (−0.5%) for negative ΔBOLD, but strong positive ΔCBF (5.9%) for positive ΔBOLD, despite similar ΔBOLD amplitudes (−0.29% and 0.32%). For this voxel subset, discordant negative ΔBOLD is unlikely due to artifacts, as a canonical hemodynamic response was confirmed for positive BOLD signal changes.

In summary, we identified a canonical hemodynamic response for mean positive ΔBOLD, but inconsistencies in certain regions. For negative ΔBOLD, we did not find any significant negative ΔCBF or ΔCMRO_2_. Our findings align with previous animal studies indicating inconsistent hemodynamic responses in cortical and subcortical regions ^31,34,63^.

### Validation of discordant hemodynamics using the Davis model and replication data

On the voxel level, we evaluated our findings in comparison to a well-established model of cerebral hemodynamic responses (Fig. 3). The Davis model predicts BOLD signal amplitudes for realistic ranges of ΔCMRO_2_ and ΔCBF in the human brain ^48^. Despite being published over 20 years ago, no study has examined the model’s accuracy across the human cortex.

Intriguingly, the Davis model predicts a range of discordant positive and negative ΔBOLD for biologically plausible ΔCBF in relation to ΔCMRO_2_ (Fig. 3, violet shadings and voxels). Our empirical findings (Fig. 3B) align closely with these predictions using established model parameters from the literature. For all voxels with concordant positive and negative ΔBOLD, we observed a canonical hemodynamic response with average *n*-ratios of 2.0 and 1.6, respectively (Fig. 3D). Our data also confirm the presence of a substantial amount of discordant voxels, comprising 31% of voxels with positive and 66% of voxels with negative ΔBOLD for CALC vs. CTRL (Fig. 3C), and similarly for the REST (Fig. S4A) and the MEM condition (Fig. S4B).

Additionally, we re-calculated ΔCMRO_2_ via the Davis model using parameters (α, M) derived from our own data, yielding a similar percentage and regional distribution of discordant voxels, along with comparable CMRO_2_ responses (Fig. S7). In conclusion, these results strengthen the reliability of the mqBOLD approach, successfully validating the decades-old model parameters of the Davis model.

One could argue that discordant ΔBOLD arises from misalignment and partial volume effects when integrating voxel data across modalities. Thus, we performed a replication study with matched voxel sizes and matrices in BOLD- and mqBOLD sequences, albeit sacrificing whole-brain coverage for higher voxel resolution in pCASL imaging. Utilizing the same study design and analysis pipeline of the main study, we again found an *n-*ratio > 1 for concordant voxels (BOLDpos/neg: 1.7/1.2) and, critically, a considerable proportion of discordant voxels (BOLDpos/neg: 40%/40%) (Fig. S6). One may also discuss sensitivity issues of mqBOLD across cortical voxels. For instance, low pCASL sensitivity or high vascular effects (high R2’ or CBV) might lead to low ΔCBF and thus artificially induce discordant ΔBOLD. Consequently, we validated our results using control masks focusing on voxels with significant ΔCBF or Δ CMRO_2_, while excluding those with high vascular contributions (Fig. S8). All control analyses confirmed the presence of discordant voxels, with both positive (11-29%) and negative (68-78%) ΔBOLD values. As a final control against voxel-specific artifacts, we investigated hemodynamic and metabolic responses for both signs of ΔBOLD within identical voxels (Fig. 4A, conjunction mask). We assessed response patterns in approximately 4.000 voxels and classified them by response type. The ‘concordant only’ voxels displayed a canonical hemodynamic response for both positive and negative ΔBOLD, demonstrating that the mqBOLD method reliably detects canonical responses across the ΔBOLD spectrum. The largest group, labeled ‘mixed’ voxels, showed a canonical hemodynamic response for positive ΔBOLD (Fig. 4B, red) but lacked a significant CBF response for negative ΔBOLD (Fig. 4B, orange). Paradoxically, interpretation based solely on BOLD fMRI data would suggest increased activity in one condition and decreased in another, despite both tasks indicating a significant increase in Δ CMRO_2_ of similar magnitude in those voxels (Fig. 4B red/orange bar plots).

Discordant voxels occurred for both positive and negative ΔBOLD, as well as for ΔCMRO_2_ derived from Fick’s formula and the Davis model. The presence of both concordant and discordant responses within identical voxels implies different hemodynamic mechanisms serving varying oxygen demands. Shaw et al. identified microvascular variations between cortical and subcortical regions ^31^ and simultaneous recordings of neuronal activity and CBF suggest that interneuron activity may influence blood flow more than other neuronal activity ^64–66^. In humans, Devi et al. ^67^ and Mullinger et al. ^16^ both found discrepancies in the coupling of CBF and BOLD signal changes in sensory cortices, which they interpreted as distinct neurovascular coupling mechanisms, particularly for negative ΔBOLD. In conclusion, mqBOLD effectively identifies canonical hemodynamic responses but also a considerable number of voxels exhibiting discordant ΔBOLD across various tasks and in relation to varying baseline conditions.

### Baseline OEF predicts baseline metabolism and alternate hemodynamic coupling

According to Fick’s principle, a change in CMRO_2_ without CBF alteration arises from ΔOEF. We hypothesized that baseline OEF varies across voxels with different hemodynamic response types, suggesting the presence of different oxygen buffers. Our findings confirmed significantly different baseline OEFs across three response types among ‘conjunction voxels’ (Fig. 4C). Baseline OEF was lowest in ‘discordant only’ voxels and highest in ‘concordant only’ voxels, with ‘mixed’ voxels in between. We also observed a significant positive linear relationship between baseline CMRO_2_, CBF, and OEF across voxels of all tasks, covering over 50% of all GM voxels (Fig. 5A, scatter plots). Multiple linear regression showed that OEF accounts for the majority of baseline CMRO_2_ variability (>68%), followed by CBF (>28%) and CBV (>1%), which also aligned with quantitative susceptibility mapping data ^68,69^ (Fig. 5C). Collectively, our results suggest that OEF is a key modulator of baseline metabolism across the human cortex, potentially predicting regional hemodynamic responses during tasks (Fig. 5D). In line with baseline results, multiple linear regression revealed that ΔOEF significantly contributes (58% of the total explained variance) to task-related oxygen demand in discordant voxels. Conversely, concordant voxels primarily accommodate ΔCMRO_2_ through ΔCBF (87%), supporting the canonical hemodynamic response. In summary, OEF is a strong predictor of baseline metabolism across the cortex and ΔOEF regulates oxygen demand in spatially distributed voxels. Our findings align with rodent studies demonstrating varying neurovascular coupling based on variations in vascular composition ^14,28,31^.

### Limitations of mqBOLD and control analyses

The mqBOLD method, while less invasive than PET, has notable limitations. Still, systematic biases in CBF, OEF, and CMRO_2_ quantification affect only across-subject comparisons and do not account for task effects, which are derived from ΔOEF or ΔCMRO_2_ within identical voxels.

Absolute quantification of CBV through DSC MRI is challenging, reflecting total rather than venous CBV. Yet, this limitation applies to nearly all available CBV measurements, including PET^70^. To enhance intersubject comparability of CBV values, we implemented a global normalization procedure ^44^. Despite these limitations, our CBV data are valuable as they represent a rare attempt to quantify baseline CBV and ΔCBV during task processing in healthy individuals.

We further evaluated the sensitivity of our mqBOLD method to accurately detect both positive and negative ΔCBF. Our analysis replicated the amplitude and extent of reduced CBF observed in a prior PET study (Fig. S3B) using a similar analysis approach ^71^. Additionally, our findings reveal ΔCBF in regions exhibiting significant positive ΔBOLD, aligning with earlier MR-based CBF measurements ^50,72^.

Furthermore, CMRO_2_ values in white matter (WM) are not interpretable since T2 and T2* quantification are influenced by orientation effects of myelinated nerve fibers and differences in lipid concentration between GM and WM. In prior work, our group addressed these systematic errors of background magnetic fields ^44^, T2-related bias ^45^, and orientation-related effects in WM structures ^73^. Additionally, we used masking to reduce partial volume effects of adjacent WM structures, confirmed our main result in a replication study, and demonstrated that mqBOLD-based OEF agreed well with PET-based OEF ^46^.

We also performed several control analyses to confirm the study design. A separate control study validated the stability of the BOLD (Fig. S3D) and CBF (Fig. S3E) signals during extended measurements. Crucially, neither signal exhibited habituation or drift effects during prolonged acquisition.

### Future directions for BOLD fMRI

Our study reveals spatial variability in hemodynamic changes across the human cortex, suggesting diverse underlying mechanisms. Firstly, voxels primarily governed by ΔOEF exhibit a greater oxygen buffer, maintaining adequate oxygen pressure during tasks ^7,19^. Secondly, OEF regulation may indicate different signaling mechanisms ^6^, including astrocytic activity ^74^, shifts in excitatory/inhibitory signaling ^75,76^, or neuromodulatory regulation ^77^. Thirdly, our findings underscore the importance of quantitative mqBOLD fMRI for future research, particularly when examining groups with altered hemodynamics, such as in aging ^30^ or neurodegenerative conditions ^78^.

## Acknowledgements

We thank Antonia Bose, André Hechler, Mahnaz Ashrafi, Roman Belenya, Laura Fraticelli, Stephan Kaczmarz, Jan Kufer and Gabriel Hoffmann for their methodological support, discussions, and suggestions on this work, and Claus Zimmer for his overall support of the study. VR acknowledges support from the European Research Council (ERC) under the European Union’s Horizon 2020 research and innovation program (ERC Starting Grant, ID 759659). CP has been funded by the Deutsche Forschungsgemeinschaft (DFG, German Research Foundation, ID 395030489).

## Author contributions statement

SE designed the study, acquired the data, performed all analyses, and wrote the manuscript

GC assisted with data analyses

BY performed Davis model analysis

JAH designed the study

CP designed the study, and developed the mqBOLD analysis pipeline

VR designed the study, supervised the project, and wrote the manuscript

## Competing Interests Statement

The authors declare no competing interests.

## METHODS

### Participants

#### Main study

Forty-seven healthy adults were enrolled; seven were excluded due to task difficulties (n=1), contrast-agent issues (n=2), or poor data quality (n=4; e.g., motion or susceptibility artifacts, unilateral CBF), leaving 40 right-handed participants (22 women, 18 men; mean age 32.1 ± 9.2 years) for analysis. Of these, 30 completed all four conditions (CALC, MEM, CTRL, REST), while 10 performed only CALC and CTRL but underwent two DSC scans, enabling within-subject ΔCBV estimation; analyses of MEM or REST thus include 30 participants.

#### Control study

To assess BOLD fMRI stability during prolonged tasks, a separate cohort of 18 right-handed healthy adults (11 women, 7 men; mean age 28.1 ± 4.8 years) underwent BOLD fMRI.

#### Replication study

For a replication with harmonized voxel matrices and resolution across BOLD and mqBOLD sequences, 10 healthy adults (5 women, 5 men; mean age 31.8 ± 6.8 years) performed CALC and CTRL.

All participants gave written informed consent. Procedures were approved by the Ethics Review Board of the Klinikum Rechts der Isar, Technical University of Munich, and participants were compensated.

#### Task design

Participants were scanned supine, viewing instructions on a screen via a mirror mounted on the head coil. Right-hand responses were recorded with a button box (Cambridge Research Systems). Tasks were explained and briefly practiced before scanning, emphasizing accuracy over speed. All tasks were designed to maintain continuous engagement during BOLD and qunatitative fMRI.

**CALC** (calculation task): This task aimed to elicit negative BOLD responses in the default mode network (DMN) and positive BOLD responses in task-positive networks. Participants solved arithmetic problems at their own pace, with a maximum response time of 10 seconds per task, following a design similar to that of Lin and colleagues ^80^. Each trial presented a row of three numbers plus a question mark (n1, n2, n3, ?) with instructions to solve the arithmetic and fill in the missing number. The solution followed the rule: n2 - n1 = DIFF, n2 = n1 + (1 * DIFF), n3 = n2 + (2 * DIFF), and= n3 + (3 * DIFF). For example, for the row 33 38 48 ?, DIFF = 5, making the correct answer 63. Participants selected from three response options, including the correct answer, each corresponding to a button on the response box.

**MEM** (autobiographical memory task): The MEM condition was based on the design of Spreng and colleagues ^53,55^, using cue words instead of pictures for consistency across tasks. Participants recalled a specific autobiographical event with as many details as possible, with the cue word displayed for up to 15 seconds. They pressed the first button twice upon recalling an event. If they couldn’t recall any details, they pressed the second and third buttons in succession for a new cue word, ensuring uniformity in button presses across tasks.

**CTRL** (low-level baseline): The CTRL condition involved a simple task with minimal cognitive demands. A row of random letters was displayed for 5.9 to 8.9 seconds, and participants pressed a button to indicate if the first letter was a vowel. This active baseline ensured visual input and button presses were consistent with the CALC and MEM tasks.

**REST** (resting state baseline): While studies on DMN activations typically use matched control tasks for contrasts ^53–55^, such as our CTRL condition, DMN deactivation studies often compare to an uncontrolled resting state baseline ^51,56,80^. To replicate these DMN contrasts, we also collected REST data, featuring a white fixation cross on a black screen.

#### MRI acquisition parameters

**Main study**: MRI was conducted on a 3T Philips Ingenia MR scanner with an Elition upgrade and a 32-channel head coil. The quantitative fMRI protocol included multi-parametric, quantitative BOLD (mqBOLD) and ASL imaging:

<u>Multi-echo spin-echo T2 mapping</u>: 3D gradient spin echo (GRASE) readout as developed by our group (Kaczmarz et al. 2020) with 8 echoes of even-spaced echo times (TE): TE1 = ΔTE = 16ms; TR=251; readout duration = 128ms; α=90°; voxel size 2×2×3.3mm^3^; 35 slices (30 slices in 4 subjects); total acquisition time = 2:28min (for 35 slices).
<u>Multi-echo gradient-echo T2* mapping,</u> as developed by our group ^44,45^: 12 echoes, TE1 = ΔTE = 5ms; TR=2229ms; readout duration = 60ms; α=30°; voxel size 2×2×3mm^3^; gap 0.3mm; 35 slices (30 slices in 4 subjects). Correction for magnetic background gradients with a standard sinc-Gauss excitation pulse ^81,82^; acquisition of half- and quarter-resolution data in k-space center for motion correction ^83^; total acquisition time = 6:08min (for 35 slices).
<u>Dynamic susceptibility imaging (DSC),</u> as described previously by our group ^84^: Single-shot GRE-EPI; EPI factor 49; 80 dynamics; TR = 2.0s; α=60°; acquisition voxel size 2×2×3.5mm^3^; 35 slices (30 slices in 4 subjects). Injection of gadolinium-based contrast agent as a bolus after 5 dynamics, 0.1ml/kg, minimum 6ml, maximum 8ml per injection, flow rate: 4ml/s, additionally flushing with 25ml NaCl; total acquisition time = 2:49min (for 35 slices).
<u>Pseudo-continuous arterial spin labeling (pCASL):</u> following Alsop et al. (2015) and as implemented by our group ^45,86^. Post-labeling delay 1800ms, label duration 1800ms; 4 background suppression pulses; 2D EPI readout; TE=11ms; TR=4500ms; α=90°; 20 slices (16 slices in one subject); EPI factor 29; acquisition voxel size 3.28×3.5×6.0mm^3^; gap 0.6mm; 39 dynamics plus one proton density-weighted M0 scan; total acquisition time = 6:00min. In addition to quantitative fMRI, we acquired
<u>BOLD-fMRI</u> using single-shot EPI, EPI factor 43; voxel size = 3.0×3.0×3.0mm^3^; FOV 192×192×127.8mm^3^; TE=30ms; TR=1.2s; α=70°; 40 slices; SENSE-factor = 2; MB-SENSE-factor = 2. 400 dynamics (8:05 min) for four conditions, 200 dynamics (4:05 min) for two conditions, 1650 dynamics (33:05 min) for the long-block control study acquisition. For susceptibility correction, a <u>B0 field map</u> was acquired with two echoes; TR/TE1/TE2=525ms/6.0ms/9.8ms; 40 slices; parallel acquisition; α=60°; voxel size = 3.0×3.0.3.0mm^3^; FOV 192×192×127.8mm^3^; total acquisition time: 0:35s.
<u>T1-weighted 3D MPRAGE</u> pre- and post-gadolinium (TI/TR/TE/α = 100ms/9ms/4ms/8°; CS-SENSE-factor=7.5; 170 slices; FOV=240x252x170 mm^3^; voxel size 1.0x1.0x1.0mm^3^; acquisition time=2:05min) and <u>T2-weighted 3D FLAIR</u> (fluid-attenuated inversion recovery; TR/TE/α = 4800/293/40°; CS-SENSE-factor=10; 140 slices; FOV=240×248.9×168mm^3^; acquisition voxel size 1.2×1.2×1.2mm^3^; turbo spin-echo factor 170; inversion delay 1650ms; acquisition time=2:09min) images were acquired for anatomical reference and to exclude brain lesions.

**Control study**: MRI was performed on a 3T Philips Ingenia scanner with a 32-channel head coil and included only T1-weighted 3D MPRAGE and BOLD-fMRI, with all acquisition parameters being identical to those described above.

**Replication study**: MRI was performed identical to the Main study, but with matching voxel matrices: T1-weighted 3D MPRAGE, multi-echo spin-echo T2 mapping and multi-echo gradient-echo T2* mapping as described above, with T2/T2* voxel size = 2×2×3.3mm^3^. <u>BOLD-</u> <u>fMRI</u> as described above, yet with a voxel acquisition size = 4x4x3mm^3^, gap 0.3mm, 40 slices. <u>PCASL</u> as described above, with BOLD fMRI voxel size (4×4×3mm^3^, gap 0.3mm; 26 slices). As a result, subject data were acquired with harmonized voxel dimensions, i.e. matching BOLD and pCASL voxel sizes and four T2/T2* voxels per BOLD/pCASL voxels in the x/y-plane. Whole-brain coverage was not possible with this higher-resolution pCASL sequence; thus, we positioned the volume at the same angle as before to cover all key regions of interest from our main study (see ’brain coverage,’ Fig. S6).

#### Data acquisition

After obtaining informed written consent, a physician placed a venous catheter for blood sampling (hemoglobin, hematocrit (Hct), creatinine). DSC contrast was administered only if creatinine was normal, ensuring renal health. Arterial oxygen saturation was tracked via pulse oximetry (Nonin Medical B.V., The Netherlands).

**Main study**: Figure 1B illustrates the imaging session from the main study.

<u>BOLD fMRI</u>: The four task conditions were presented using a 30-second block design, each repeated four times in random order. BOLD alternated with mqBOLD fMRI runs to reduce habituation or fatigue effects. During mqBOLD imaging, conditions were presented pseudo-randomly for pCASL and T2* mapping. DSC (CTRL) and T1w post-gadolinium scans were performed at the end of the session to avoid signal artifacts. The contrast agent was administered via a pump (Medtron AG, Saarbrücken, Germany) under a medical doctor’s supervision.

After the imaging session, participants completed a memory questionnaire about the MEM condition, rating the ease of recalling specific events on a difficulty scale (1 = ’very easy’ to 4 = ’very difficult’) and the detail of their memories on a concreteness scale (1 = ’very detailed’ to 4 = ’very vague’). On average, participants scored 1.8 ± 0.7 on difficulty and 2.0 ± 0.6 on concreteness, indicating that recalling events was relatively easy and memories were reasonably detailed. Participants took an average of only 2.5 ± 1 seconds to recall an event.

A subsample of N=10 subjects from the main study performed only the CALC and CTRL conditions but instead received two DSC scans (totaling a full clinical dose of 16ml). This enabled calculation of within-subject ΔCBV for task effects.

**Control study**: fMRI BOLD data were obtained for CALC and MEM tasks with a 30s block design (4 repetitions each, interleaved with 30s CTRL blocks), alongside extended 3-minute blocks (4 repetitions each, interleaved with 1min CTRL blocks), totaling 41 minutes of scan time.

**Replication study**: Study design and data acquisition were identical to the main study, utilizing the group median CBV data from the main study for analyses.

##### Image processing

### Processing of BOLD fMRI data

BOLD fMRI data were pre-processed using fMRIPrep 20.2.4 ^87^ within a docker container based on Nipype 1.6.1 ^88^. Preprocessing involved: segmentation, estimation of motion parameters and other confounds, correction for susceptibility distortions, co-registration in native T1w space, and normalization to MNI152-ICBM-2mm space with a non-linear 6^th^ generation registration model developed by (Montreal Neurological Institute, McGill University). fMRIPrep relies on FSL 5.0.9 for registering EPI time-series data to T1w data with boundary-based registration (BBR), FSL FAST for brain tissue segmentation, and ANTs 2.3.3 ^89^ for spatial normalization to MNI space in a multiscale, mutual-information based, nonlinear registration scheme, where all transforms are first concatenated and registration steps applied at once. As part of the fMRIPrep pipeline ^90,91^, correction for head motion and susceptibility distortions was performed in subject’s native space, applying a single composite transform to the BOLD-fMRI time series. The same data were also resampled into standard space, generating a preprocessed BOLD run in MNI152NLin6Asym space. Preprocessed BOLD fMRI data, without global signal regression, were then used as input to the PLS model. All datasets from the main, control, and replication studies underwent identical processing.

### Calculation of quantitative parameter maps from mqBOLD data

We calculated quantitative parameter maps using in-house scripts (in Matlab) and SPM12 (Wellcome Trust Centre for Neuroimaging, UCL, London, UK). Fig.1 illustrates the procedure and shows representative subject-averaged parameter maps.

<u>T2/T2*-mapping</u>: Quantitative T2 and T2* parameter maps are obtained by applying mono-exponential fits to multi-echo spin and gradient echo data, as described by our group ^44,45,92^. Corrections for macroscopic magnetic background fields were implemented ^82^ and motion artifacts were addressed through redundant acquisitions of k-space centers ^83^.

#### <u>R2’ maps</u> are calculated via

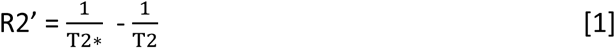

yielding the transverse, reversible relaxation rate that depends on the vascular dHb content within a voxel ^93,94^. Caution is required when interpreting these values at air-tissue boundaries (magnetic field inhomogeneities), in deep GM (iron deposition) or in WM structures (orientation effects in myelin), as previously discussed ^45,82^.

<u>CBV maps</u> are derived from DSC MRI after contrast agent application via integration of leakage-corrected ΔR2*-curves ^95^ and subsequent normalization to a white matter value of 2.5% ^96^. The DSC procedure has been described by our group ^84,97^.

#### <u>OEF maps</u> are calculated from R2’ and CBV parameter maps via the mqBOLD approach ^39,43^ and, as implemented by our group ^44^ via

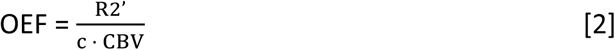

with c = γ · ^4^ · π · Δχ_0_ · *hct* · B_0_ (gyromagnetic ratio γ = 2.675 · 10^8^ s^-1^ T^-1^; susceptibility difference between fully deoxygenated and oxygenated hemoglobin Δχ_0_ = 0.264 · 10^-6^; magnetic field strength B_0_ = 3T; small-vessel hematocrit *hct*, calculated as 85% of the (large-vessel) hematocrit values measured in each subject ^44,98^). OEF (ratio) represents the amount of oxygen extracted from capillaries among passage.

<u>CBF maps</u> were calculated from pCASL data as described in ^85^. Specifically, CBF is calculated as the pairwise difference of the averaged and motion-corrected label and control images, and then scaled by a proton-density-weighted image.

CMRO2 maps: For each condition separately, we calculated the voxel-wise CMRO2 by combining all parameter maps via Fick’s principle:

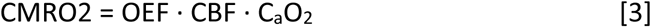

where C_a_O_2_ reflects the arterial oxygen content (in µmol) for each subject and was calculated as C_a_O_2_ = 0.334 · Hct · 55.6 · O_2_sat, with O_2_sat being the oxygen saturation measured by a pulse oximeter ^94^.

All parameter maps were calculated in subject space and co-registered to the first echo of the T2 data. Any normalization into MNI space was performed afterwards. Specifically, parameter maps were first co-registered to native T1w space before applying the normalization matrix to MNI space, as derived from fMRIPrep. CBF values were upscaled by 25% to account for the systematic underestimation of CBF due to the four background suppression pulses, as motivated here ^99,100^. The data of the replication study were processed identically to those of the main study.

### Artifact correction and GM masking

For our analyses in standard space, we excluded voxels that fell within the lowest 15th percentile of the temporal signal-to-noise ratio (tSNR) for over 66% of participants, based on the BOLD fMRI data from each subject and voxel. The excluded voxels were primarily found in regions with significant susceptibility artifacts, such as the fronto- and temporo-basal brain areas. Additionally, we masked out the cerebellum and any voxels with a GM probability of less than 0.5. The resulting group mask was then applied to both the output of the GLM group analysis and the input matrices for the partial least squares analyses. For the analyses in native space, we additionally masked CSF-prone areas (T2 > 90ms), high-susceptibility areas (R2’ > 9 s^-1^), voxels with a high percentage of blood volume (CBV >10%, probably driven by larger veins/arteries) and voxels with biologically implausible values, such as T2’ > 90ms, OEF > 0.9, and CBF > 90ml/100g/min.

### Estimation of a realistic surrogate for CBV during CALC

After a sensitivity analysis at our institution, we found that half the clinical dosage of contrast agent was sufficient to reliably assess CBV in healthy subjects. Therefore, the final subset of 10 participants received two half dosages during the CALC and CTRL conditions, allowing us to quantify CBV in both conditions. We calculated the voxel ΔCBV for CALC compared to CTRL in subject space and averaged the results across Glasser’s 360 functional ROIs ^101^. This subject-averaged ΔCBV map was then used to estimate a CBV surrogate map for the remaining 30 subjects, who only had one baseline CBV measurement. To our knowledge, this is the most empirically supported data for ΔCBV in quantitative fMRI studies, but it was only possible for the CALC condition. Hence, we continued using the CBV CTRL map to calculate CMRO2 during the MEM condition.

### Semi-quantitative, BOLD-informed, CMRO2 estimation

The estimation of quantitative CMRO2 maps relies on the combination of R2’, CBV, and CBF values, according to Eq. [3]. To control for potential error propagation during voxel-wise analyses, especially from R2’ measurements (see appendix in ^73^, we calculated R2’ parameter maps during MEM and CALC from baseline R2’- and BOLD fMRI data as suggested by Fujita et al. ^102^. Instead of calculating ΔR2’ from quantitative R2’ (multi-echo GE-based T2*) during task conditions, ΔR2’ is approximated as

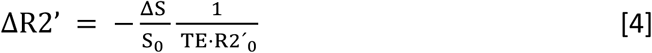

with TE = 30ms, R2’_0_ being the baseline R2’, and ^ΔS^ the BOLD signal change, derived from task S_0_ data. R2’ in CALC and MEM was calculated via R2’ = R2’_0_ + (ΔR2’ · R2’_0_) and fed into OEF and CMRO2 calculations according to Eqs. [2] and [3]. These semi-quantitative CMRO2 parameter maps differ from the regional CMRO2 maps only in their underlying R2’ values. In Fig. S5, we compared the PLS results of the BOLD-informed approach with those of the fully quantitative approach and found very similar signal ranges and voxel distributions.

### Davis model

The Davis model was originally designed for calibrated fMRI and simulates ΔBOLD (= ΔS/S_0_) by using carbon dioxide breathing as a physiological method to manipulate CBF independently of CMRO2. The model relies on parameters M, α, and β ^48^:

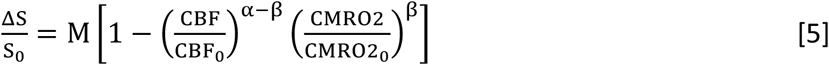

where α is a power-law exponent relating ΔCBV to ΔCBF, β reflects the microvascular anatomy ^94^ and M is commonly referred to the maximum ΔBOLD that occurs when all dHb is removed from the vessels and depends on baseline OEF ^93^. Previously, α = 0.38 was used ^103^, but more recent studies have identified lower values. In Fig. 3A, we plotted the range of predicted ΔBOLD using empirical values for all parameters, derived from recent calibration studies with α = 0.23, β = 1.3 ^94^, and M=5.5 ^78^. Assuming α and β, the fractional change in CMRO2 can be calculated from combined BOLD signal and CBF measurements during tasks ^78,93,104^.

### Statistics

#### Partial least squares analysis of BOLD and mqBOLD data

Partial least squares (PLS) analyses were performed using the pyls library in Python language (Python Software Foundation, version 3.8). Mean-centered PLS is a data-reduction method that computes latent variables (LVs) and corresponding brain patterns, optimizing the relationship between brain signals and experimental design ^105^. In this study, we used PLS analyses to perform group-level statistics to identify brain regions that distinguish between task conditions (CALC or MEM) and a baseline condition (CTRL or REST). This analysis was applied to both BOLD fMRI and mqBOLD data, allowing for comparison of statistical maps. For mqBOLD data, we used quantitative values OEF, CBF, or CMRO2 values per voxel and subject.

For BOLD fMRI data, we used median percent signal change (from either CTRL or REST) across 24s (20 TRs) per task condition, excluding the first 6s of each task block to account for the hemodynamic response lag.

The significance of the LVs (multivariate patterns) was tested using permutation tests (3000 permutations), while the reliability of brain saliences (voxel contributions) was assessed through bootstrap resampling (3000 samples). Significant brain regions were identified using the bootstrap ratio (BSR), where a BSR > ±2 suggests a 95% confidence interval if the bootstrap distribution is normal ^106,107^. To interpret BSR maps, they must be compared to design scores of each condition within each LV for task differences. In this manuscript, orange-red indicates a positive task effect (higher values during the task versus baseline), while blue indicates the opposite. We visualized the whole-brain, non-thresholded PLS results by transforming statistical maps from native to surface space with the “vol_to_surf” function from nilearn’s surface toolbox, and plotted them on the “fsaverage pial left and right mesh” using “plot_surf_stat_mat.”

### BOLD clusters

The BOLD clusters in Fig. 2D are based on a PLS analysis of BOLD data, with statistical maps thresholded at a BSR score of >+/-3. We extracted regions with a size of >1,000 connected voxels using Nilearn’s region package in Python and fused fragmented clusters that belonged to the same Yeo’s network upon visual inspection ^79^.

### General linear modeling of BOLD data

To validate the PLS analyses, we used a General Linear Model (GLM) approach as recommended by ^108^. The GLM included confound variables such as CSF and WM signals, dvars, framewise displacement, and translations/rotations across x-, y-, and z-axes. We applied a high-pass filter (100s) and a 6 mm smoothing kernel. For native space analyses, we used individual first-level z-maps (z > 2.5). The contrasts calculated were: CALC-positive (CALC > CTRL or REST), CALC-negative (CALC < CTRL or REST), MEM-positive (MEM > CTRL), and MEM-negative (MEM < CTRL). To cross-validate the PLS results, we used FWE-corrected z-maps from the second-level analysis.

### Other statistical analyses

For the native space analyses, we calculated median values within each native-space ROI from the first-level GLM output (z > 2.5) for each subject. We assessed significant task-related differences compared to baseline using paired-samples two-sided t-tests across subjects. For other analyses, we obtained median voxel values in standard space across subjects (see Figs. 2B, 3B, and 4A&D). Bar plots (Fig. 2D) were created using Python’s seaborn library ^109^ with error bars indicating a 95% confidence interval (CI) based on 2000 bootstraps. A CI that included zero indicated a statistically non-significant median delta value. To analyze hemodynamic differences between discordant and concordant voxels, we conducted paired t-tests on median subject values, while baseline differences were assessed using independent-sample permutation tests. High-resolution external maps were downsampled to 2mm MNI standard space prior to analysis.

## DATA AVAILABILITY

All raw and processed data are publicly available on OpenNEURO ^110^ https://openneuro.org/datasets/ds004873.

## CODE AVAILABILITY

The scripts and Python Jupyter notebooks for quantitative parameter map analyses, and the configuration files for replication of all analyses and figures are available on GitHub: https://github.com/NeuroenergeticsLab/two_modes_of_hemodynamics.

The scripts for generating the parameter maps are available on GitHub: https://gitlab.lrz.de/nmrm_lab/public_projects/mq-bold.

## Notes

https://openneuro.org/datasets/ds004873

### Competing Interest Statement

The authors have declared no competing interest.

### Summary of Updates

This is the accepted version of the manuscript. We have changed the title and have significantly shortened both the abstract and the manuscript. No significant changes have been made content-wise.

