## Supplementary material for "BOLD signal changes can oppose oxygen metabolism across the human cortex": Fig_S1_to_S8

**by Epp et al.**

**METHODS** (supplementary material only)

Fig. S2: To compare our data with CBF data from PET imaging, we followed the approach outlined by Shulman et al. (Shulman et al., 1997). We calculated the median ΔCBF [%] across subjects using a threshold of CBF < -2% and no further corrections (Fig. S2B). CBF peak ROIs were obtained from the PLS-derived statistical map (Fig. S2B, right). Using the atlasreader library in Python (Notter et al., 2019), we selected nine peaks with the highest positive values (out of 15) and all seven negative peaks. We created 5mm diameter spheres around these coordinates and extracted median CBF and CMRO2 responses to compare CALC and REST conditions to the original Shulman et al. figure.

Fig. S8: In a validation experiment shown in Fig. S8, we recalculated CMRO2 values using the Davis model instead of Fick's formula. We calculated α and M in native space from the CTRL data (N=40) and extracted median values for Glasser ROIs (Glasser et al., 2016), and assuming a power-law relationship between CBV and CBF: $CBV = 0.8\,CBF^{\alpha}$ Grubb et al. (1974). Fitting this to our data, with venous CBV as 0.7 times total CBV, yielded α = 0.38. We also calculated a subject-specific calibration factor M = TE * R2', where R2' was based on venous CBV, resulting in an average M of 11.2 ± 1% across 40 subjects.

**Table S1**

*Mean BOLD signal change and parameter values of the mqBOLD approach, in baseline and as task changes across subjects (standard deviation) within individual first-level-GLM BOLD-fMRI masks, separately for positive and negative BOLD responses.*

| **task vs. baseline** | **BOLD mask** | **BOLD** [%] | **CBF** [ml/100g/min] | | **CMRO2** [µmol/100g/min] | | **OEF** [ratio] | |
|---|---|---|---|---|---|---|---|---|
| | | | baseline | ΔCBF | baseline | ΔCMRO2 | baseline | ΔOEF |
| CALC vs. CTRL | CALC-pos | **0.70%** | 46.5 (8.2) | **7.7%**<br>3.1** (3.2) | 124.0 (20.9) | **4.9%**<br>6.4** (7.7) | 0.37 (0.06) | **-3.2%**<br>-0.013** (0.01) |
| | CALC-neg | **-0.34%** | 46.6 (6.4) | -1.9%<br>n.s. | 120.1(18.1) | -0.3%<br>n.s. | 0.35 (0.05) | 0.01%<br>n.s. |
| MEM vs. CTRL | MEM-pos | **0.51%** | 48.5 (9.1) | **5.6%**<br>2.6** (2.8) | 125.0 (18.8) | **5.1%**<br>6.4** (8.2) | 0.36 (0.06) | **-0.7%**<br>-0.003* (0.013) |
| | MEM-neg | **-0.35%** | 48.4 (8.9) | -1.7%<br>n.s. | 126.7 (31.1) | -1.1%<br>n.s. | 0.36 (0.07) | 1.1%<br>n.s. |
| CALC vs. REST | CALC-pos | **1.01%** | 45.3 (8.8) | **10.0%**<br>4.4** (3.2) | 123.5 (24.5) | **6.4%**<br>6.5** (8.1) | 0.38 (0.06) | **-3.6%**<br>-0.014** (-0.014) |
| | CALC-neg | **-0.38%** | 47.7 (7.8) | -2.6%<br>-1.2 (3.6) | 123.7 (20.3) | -0.9%<br>n.s. | 0.36 (0.06) | 1.0%<br>n.s. |

*Note.* Quantitative values of CBF, CMRO2, and OEF are displayed both at baseline and as absolute delta values (task minus baseline, median across 40 subjects for CALC vs. CTRL and median across 30 subjects for MEM vs. CTRL and CALC vs. REST), along with percent change values (bold font, if significant) for each contrast. Values reflect data as displayed in Fig. 2C, with CMRO2 values in CALC being corrected for CBV increases, see Methods. ** $p < .001$, based on paired, two-sided t-tests, $df = 39$ for CALC vs. CTRL contrasts, $df = 29$ for CALC vs. REST and MEM vs. CTRL contrasts.

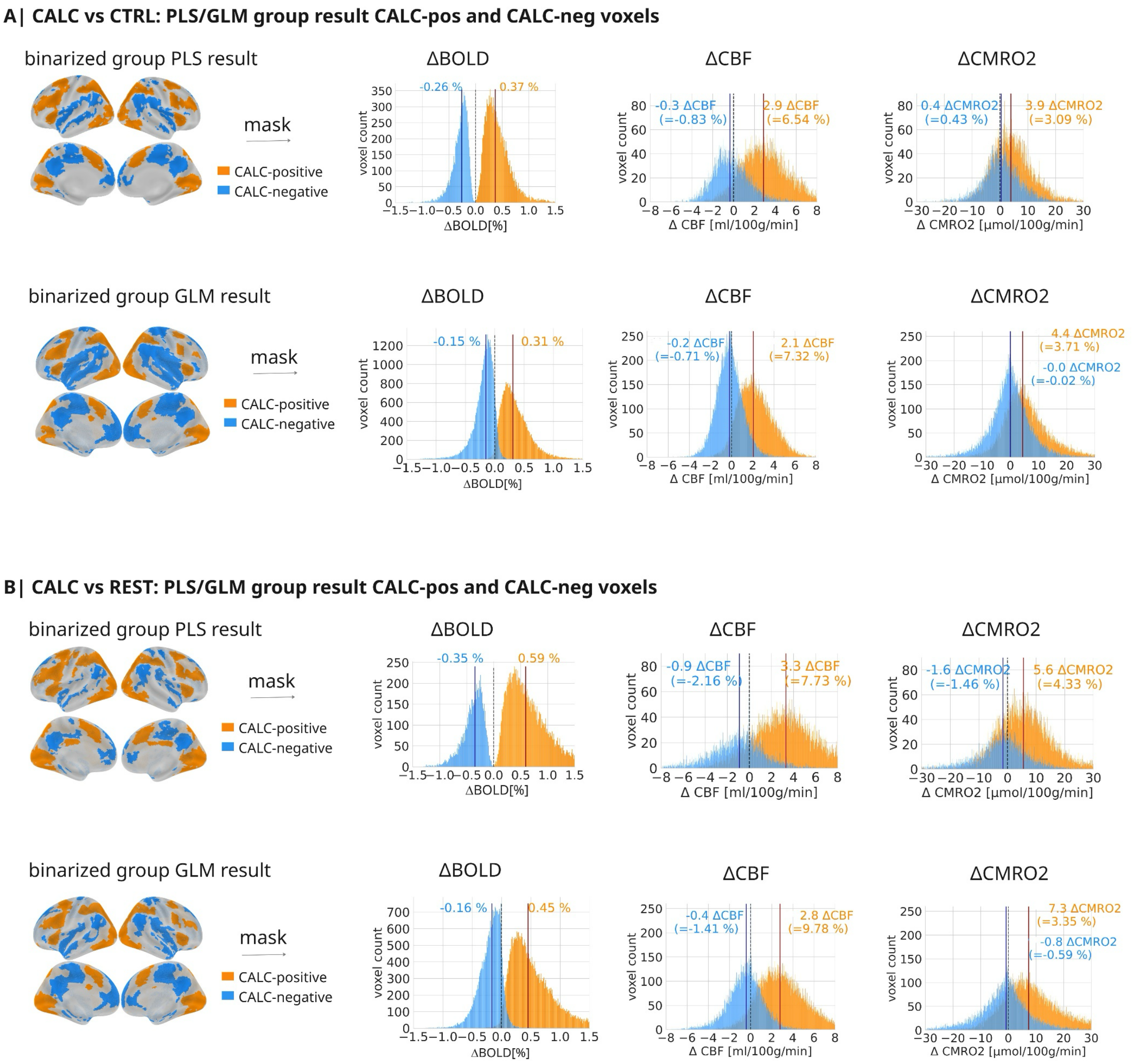


**Figure S1. Comparing ΔBOLD, ΔCBF and ΔCMRO2 in binarized positive and negative masks derived from PLS and GLM BOLD-fMRI group results, CALC vs CTRL across 40 subjects and CALC vs REST across 30 subjects. A|** CALC vs CTRL contrast, PLS results, as shown in Fig. 2 and GLM results. The CALC-negative areas are more extended compared to the PLS group results but the median BOLD responses within these areas are smaller (-0.15% vs -0.26%). Overall, results are very similar when masking with the GLM group mask vs. the PLS group mask, with CBF decreases below 1% and CMRO2 changes around zero in CALC-negative areas. **B|** CALC vs REST contrast, PLS results, as shown in Fig. S1., and GLM result. Again, the CALC-negative areas when analyzed with the GLM are more extended compared to the PLS CALC-negative areas, and the median BOLD responses within those areas is smaller (-0.16% vs. -0.35% in the PLS CALC-negative areas). Overall, the median CBF and CMRO2 responses within the CALC-negative areas are quite similar when compared within the GLM versus the PLS masks, with CBF decreases of -2.16% (PLS mask) vs. -1.41% (GLM mask and CMRO2 decreases of -1.46% (PLS mask) vs. -0.59% (GLM mask). Thus, using masks derived from GLM analysis of BOLD-fMRI instead of PLS does not change our main results and even results in weaker BOLD, CBF and CMRO2 decreases within BOLD-negative areas.

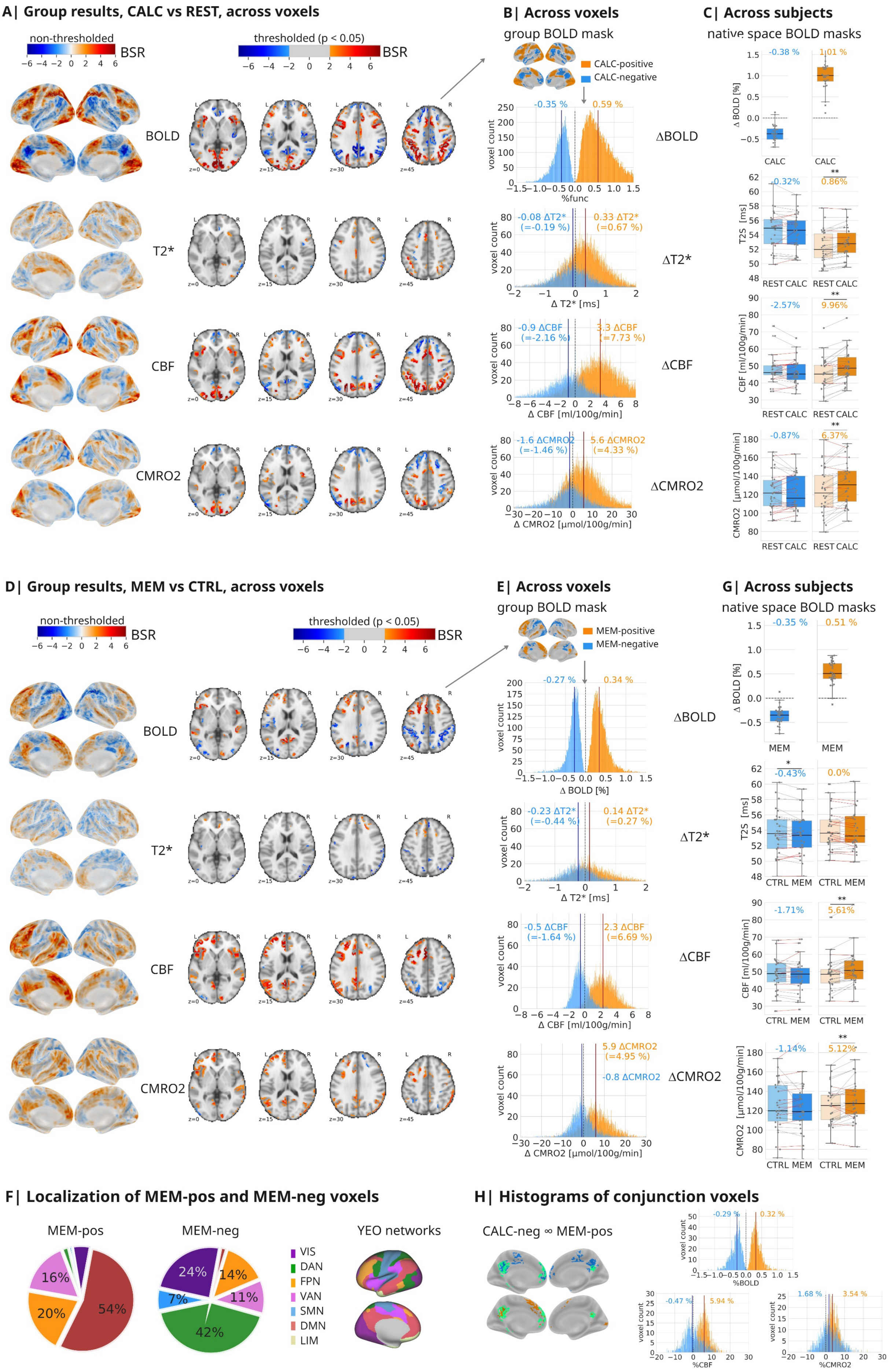

A| Group results, CALC vs REST, across voxels
non-thresholded
BSR
thresholded (p < 0.05)
BOLD
T2*
CBF
CMRO2
B| Across voxels
group BOLD mask
CALC-positive
CALC-negative
-0.35 %
0.59 %
voxel count
%func
ΔBOLD
-0.08 ΔT2*
(=-0.19 %)
0.33 ΔT2*
(=0.67 %)
Δ T2* [ms]
ΔT2*
-0.9 ΔCBF
(=-2.16 %)
3.3 ΔCBF
(=7.73 %)
Δ CBF [ml/100g/min]
ΔCBF
-1.6 ΔCMRO2
(=-1.46 %)
5.6 ΔCMRO2
(=4.33 %)
Δ CMRO2 [µmol/100g/min]
ΔCMRO2
C| Across subjects
native space BOLD masks
-0.38 %
1.01 %
Δ BOLD [%]
CALC
CALC
-0.32%
0.86%
T2S [ms]
REST CALC
REST CALC
-2.57%
9.96%
CBF [ml/100g/min]
-0.87%
6.37%
CMRO2 [µmol/100g/min]
D| Group results, MEM vs CTRL, across voxels
non-thresholded
BSR
thresholded (p < 0.05)
BOLD
T2*
CBF
CMRO2
E| Across voxels
group BOLD mask
MEM-positive
MEM-negative
-0.27 %
0.34 %
Δ BOLD [%]
ΔBOLD
-0.23 ΔT2*
(=-0.44 %)
0.14 ΔT2*
(=0.27 %)
Δ T2* [ms]
ΔT2*
-0.5 ΔCBF
(=-1.64 %)
2.3 ΔCBF
(=6.69 %)
Δ CBF [ml/100g/min]
ΔCBF
5.9 ΔCMRO2
(=4.95 %)
-0.8 ΔCMRO2
ΔCMRO2
Δ CMRO2 [µmol/100g/min]
G| Across subjects
native space BOLD masks
-0.35 %
0.51 %
MEM
MEM
-0.43%
0.0%
CTRL MEM
CTRL MEM
-1.71%
5.61%
-1.14%
5.12%
F| Localization of MEM-pos and MEM-neg voxels
MEM-pos
16%
20%
54%
MEM-neg
24%
14%
7%
11%
42%
VIS
DAN
FPN
VAN
SMN
DMN
LIM
YEO networks
H| Histograms of conjunction voxels
CALC-neg ∞ MEM-pos
-0.29 %
0.32 %
%BOLD
-0.47 %
5.94 %
%CBF
1.68 %
3.54 %
%CMRO2

**Figure S2. Replication of Figure 2 with CALC vs REST and MEM vs CTRL contrasts, across 30 subjects. A|** Statistical results of the PLS group analysis in standard space, CALC vs REST, on the BOLD data. Maps on the brain surface show non-thresholded statistical maps (values are bootstrap ratios=BSR, akin to z-values) of the first latent variable, which was significant (permutation tests, $p < .001$). Red voxels indicate higher values during CALC compared to REST, while blue voxels represent the opposite contrast. Axial slices display significant brain clusters, thresholded at BSR ±2 (akin to $p < .05$, cluster size >30 voxels). **B|** Histograms depict group-average median voxel distributions of all parameters (CALC minus REST) within the binarized CALC-positive and CALC-negative group masks (shown as surface plot). **C|** Subject distribution of ΔBOLD, ΔCBF and ΔCMRO2 within individual native-space BOLD-fMRI masks of significant positive (orange) and negative (blue) BOLD responses. Dots represent subject parameter values, calculated as the median across voxels within individual masks. Boxplots: line, median; box limits, upper and lower quartiles; whiskers, minimum and maximum data points except for outliers: values outside of 1.5* IQR; individual dots, one dot per subject, median voxel values). Paired samples t-tests, two-sided, ** = $p < .001$. Gray lines connect values obtained from individual subjects, red lines indicate subjects where the sign of ΔCBF and ΔCMRO2 deviates from the expected direction. **D|** Statistical results of the PLS group analysis in standard space, comparing MEM and CTRL, for both BOLD and quantitative fMRI parameter maps. Maps on the brain surface show non-thresholded statistical maps (values are bootstrap ratios=BSR, akin to z-values) of the first latent variable, which was significant in each analysis (permutation tests, $p < .001$). Red voxels indicate higher values during MEM compared to CTRL, while blue voxels represent the opposite contrast. Axial slices display significant brain clusters, thresholded at BSR ±2 (akin to $p < .05$, cluster size >30 voxels) for BOLD-fMRI, CBF, and CMRO2. **E|** Histograms depict median voxel distributions across subjects of ΔBOLD, ΔCBF and ΔCMRO2 (MEM minus CTRL) within the binarized MEM-pos and MEM-neg group masks (shown as surface plot). **F|** MEM-pos and MEM-neg voxels were located in different functional networks, MEM-pos primarily in the DMN and MEM-neg primarily in the VAN. Functional networks defined by Yeo et al. (2011): visual (VIS); dorsal attention (DAN); frontoparietal (FPN); ventral attention (VAN); somatomotor (SMN); default mode (DMN) network. **G|** Subject distribution of ΔBOLD, ΔCBF and ΔCMRO2 within individual native-space BOLD-fMRI masks of significant positive (orange) and negative (blue) BOLD responses. Dots represent subject parameter values, calculated as the median across voxels within individual 1st level GLM masks. Boxplots: line, median; box limits, upper and lower quartiles; whiskers, minimum and maximum data points except for outliers: values outside of 1.5* IQR; individual dots, one dot per subject, median voxel values). Paired samples t-tests, two-sided, ** = $p < .001$. Gray lines connect values obtained from individual subjects, red lines indicate subjects where the sign of ΔCBF and ΔCMRO2 deviates from the expected direction. **H|** Distribution of ΔBOLD, ΔCBF and ΔCMRO2 across conjunction voxels (displayed in green) that displayed positive ΔBOLD during MEM (orange) and negative ΔBOLD during CALC (blue), primarily located in DMN regions.

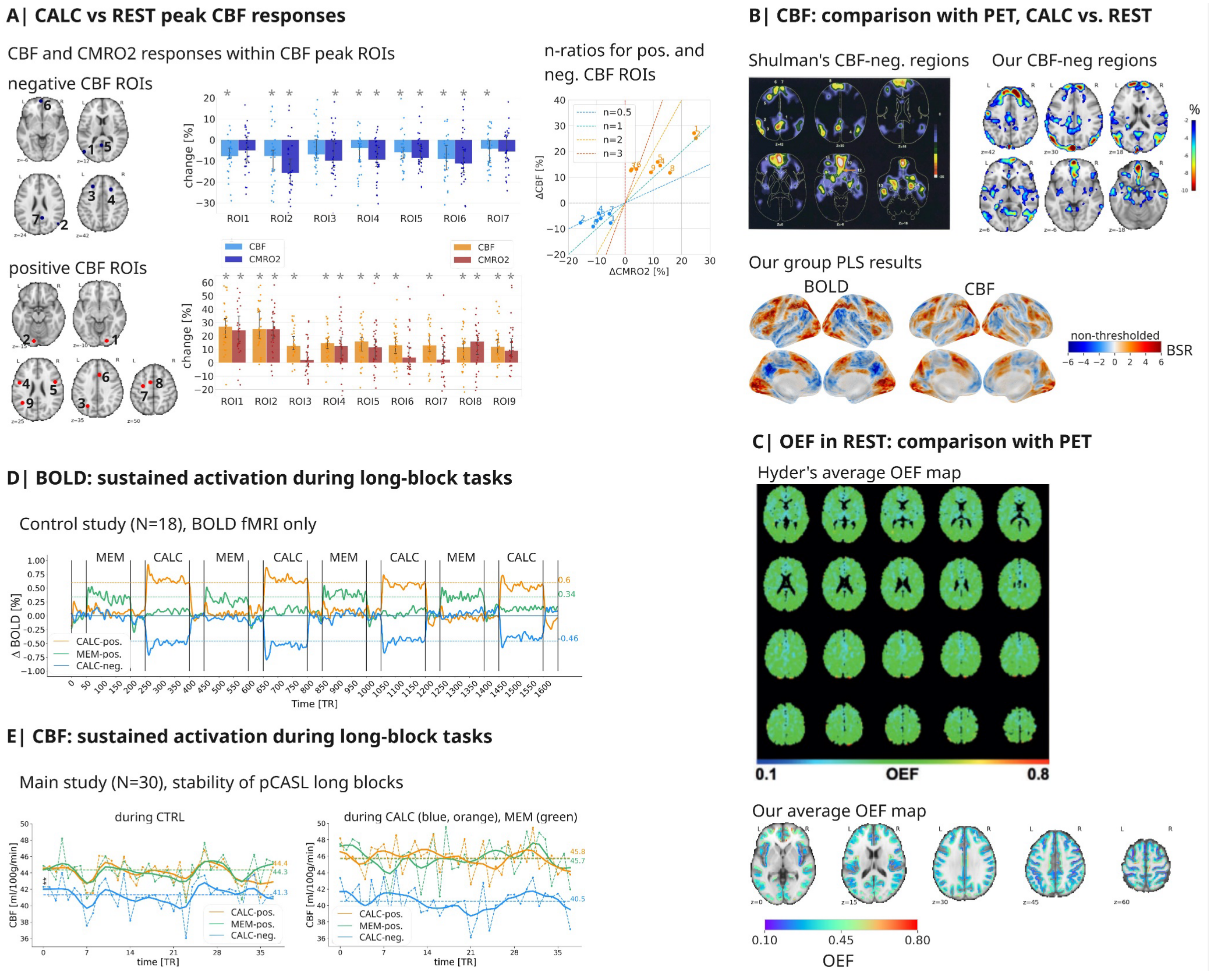


**Fig. S3. Control analyses. A|** Maximum CBF responses in CBF peak ROIs, CALC vs. REST, across 30 subjects; peaks derived from the PLS statistical CBF group map as shown in B| for CBF data, separately for positive and negative CBF changes. Bar plots show median responses within each 5mm sphere, error bars 95% CI, significant deviations from zero are denoted by asterisks (*p*<.05), not corrected for multiple comparisons. Average negative CBF responses within peak ROIs ranged from -5% to -10.5%, while positive CBF responses ranged from 12% to 29.5%. Negative CMRO2 responses within the same ROIs ranged from -2% to -10%, positive CMRO2 responses ranged from 3% to 17%. The scatterplot displays ΔCMRO2 versus ΔCBF, median per peak ROI. The lines depict *n*-ratios from 0.5 (blue) to 3 (red). *n*-ratios were calculated as the ratio between ΔCBF divided by ΔCMRO2, separately for each CBF peak ROI. **B|** We replicated Shulman et al.'s CBF-negative regions, with permission from Shulman et al. (J Cogn Neurosci, 1997), with our own CBF data, CALC vs REST. Median percent-signal change values across 30 subjects, CBF < -2%, no correction applied, showed a very similar pattern to the pooled CBF data taken from PET imaging as in Shulman et al. On the very right, statistical maps of group results, CALC vs. REST for BOLD and CBF data, non-thresholded bootstrap ratios, are shown. Please note that the negative BOLD clusters are much larger than the negative CBF clusters in DMN regions. **C|** We replicated Hyder al.'s OEF values in baseline, across 30 subjects, with permission from Hyder et al. (JCBFM, 2016), with our own OEF data in REST baseline, within gray-matter and thresholded by R2' and CBV (2.5 x absolute standard deviation around the median). **D|** Results of the BOLD-fMRI only control study. Median time-courses of BOLD fMRI signal extracted from individual activation areas derived from GLM 1st level contrast maps, z > 2.5, native space, duration task/baseline: three/one minutes, and averaged across subjects (N = 18, independent control sample). CALC > CTRL, orange; CALC < CTRL, blue; MEM > CTRL, green. BOLD signal peaked at the beginning of each task block and then stabilized at an elevated level during the entire task duration. **E|** Median CBF time-courses within CALC-positive (orange), CALC-negative (blue) and MEM-positive (green) BOLD regions, averaged across all subjects from the main study (N = 40; N = 30 for MEM). The left plot shows time-courses within the different ROIs during CTRL, and the right plot shows time-courses during CALC in CALC-neg regions in blue, in CALC-positive regions in orange and within MEM-positive regions during MEM in green. Within all conditions and ROIs, CBF was stable during 6min of acquisition.

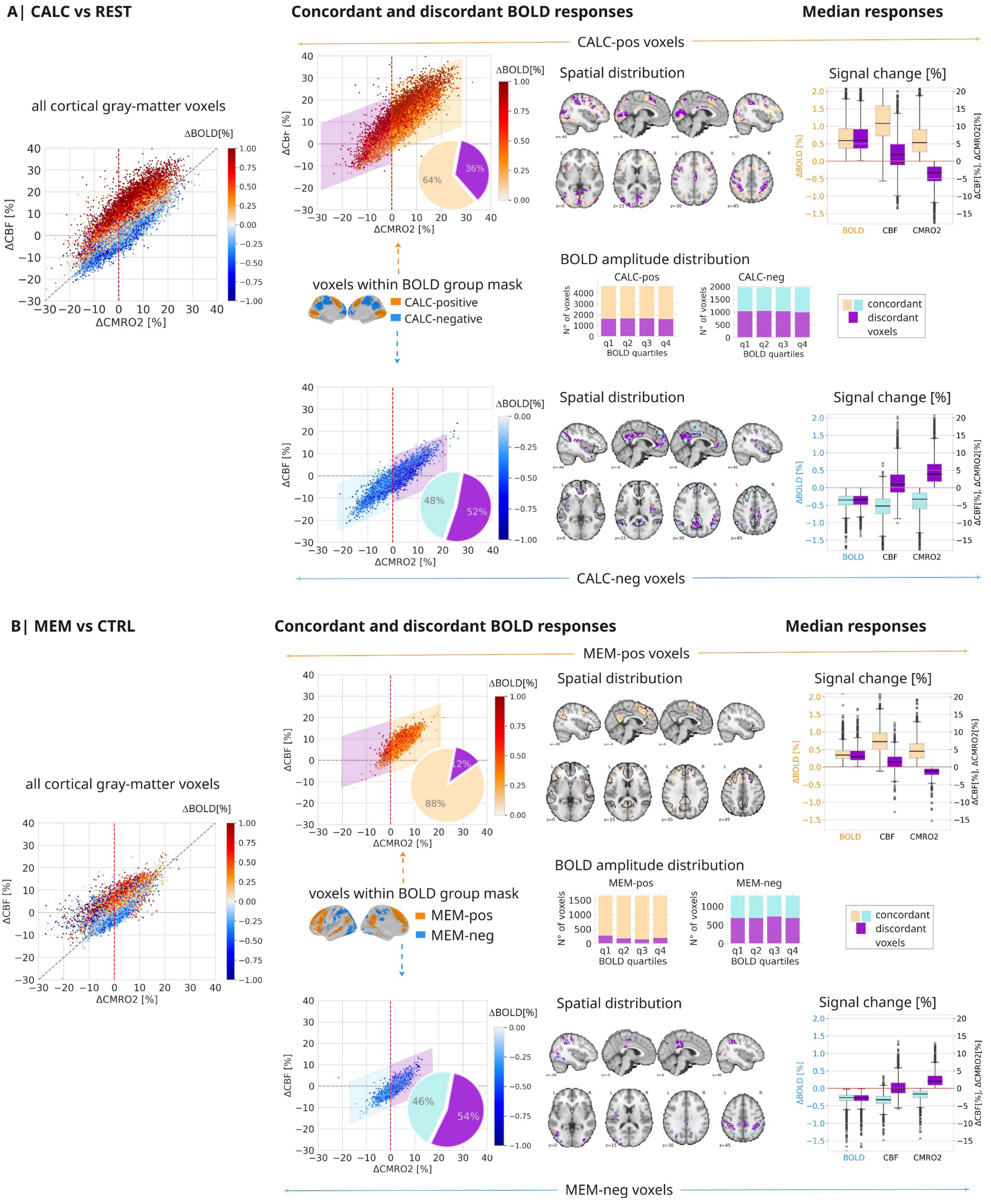


**Figure S4. Replication of Fig 3 for CALC vs REST and MEM vs CTRL, across 30 subjects. A|** CALC vs REST. Scatterplot: observed data from BOLD- and quantitative fMRI for all cortical voxels on the left, separately for significant CALC-negative and CALC-positive voxels in the middle, median across all subjects per voxel, colors indicate ΔBOLD [%]. Voxels with ΔBOLD discordant to ΔCMRO2 are highlighted in violet color. The pie plots summarize the ratio of discordant to concordant voxels. Axial slices illustrate the spatial distribution of significant concordant and discordant BOLD voxels. The stacked bars in the right center depict the amplitude distribution of discordant and concordant voxels across BOLD amplitude quartiles. It is noteworthy that discordant voxels show neither spatial nor amplitude preference. The boxplots summarize ΔCMRO2[%] and ΔCBF[%] across all voxels with either concordant or discordant ΔBOLD[%]; line, median; box limits, upper and lower quartiles; whiskers, minimum and maximum data points except for outliers: values outside of 1.5* IQR; individual dots, median voxel values across subjects. Please note that discordant and concordant voxels feature a similar ΔBOLD amplitude despite indicating opposite metabolic responses. **B|** MEM vs CTRL. Replication of A| but with MEM vs CTRL contrast.

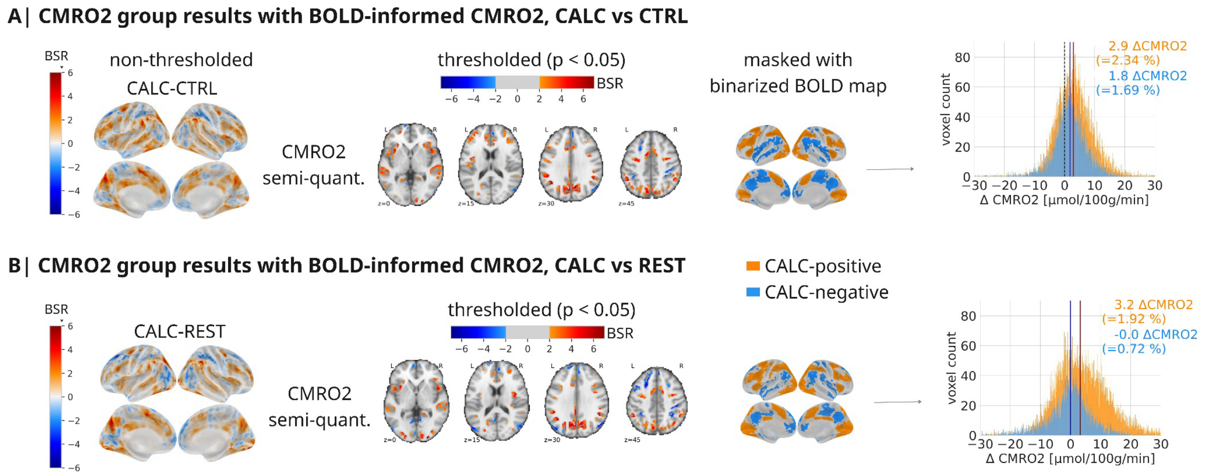


**Figure S5. PLS group analysis, CALC vs CTRL (N=40) and CALC vs REST (N=30), with BOLD-informed CMRO2.** Group results as in Fig. 2A with semi-quantitative CMRO2 data, upper row CALC vs CTRL, lower row CALC vs REST. As our goal was to explain voxel-wise changes in BOLD via changes in CBF and CMRO2, we aimed to improve the SNR of our CMRO2 task effects. Because especially T2* and therefore R2' measurements are heavily influenced by motion artifacts and depend on the correction of background magnetic field gradients, we approximated ΔR2' from BOLD fMRI data and recalculated OEF and CMRO2 task (CALC and MEM) maps, see Methods. Semi-quantitative ΔR2' and thus task CMRO2 maps benefit from motion correction and offer a higher SNR due to repeated task conditions. Our main results remained valid and were even strengthened when the main PLS group analysis was repeated using the semi-quantitative CMRO2 data. As in the full quantitative analysis, the group statistical maps showed widespread CALC activations and concordant, significant CMRO2 increases in CALC-pos BOLD regions. Yet, as before, we found no CMRO2 decreases in CALC-neg regions, independent of baseline. Compared to the full quantitative approach, CMRO2 even increased slightly, 1.7% vs. CTRL and 0.7% vs. REST (full quantitative approach: 0% and -0.6%, respectively).

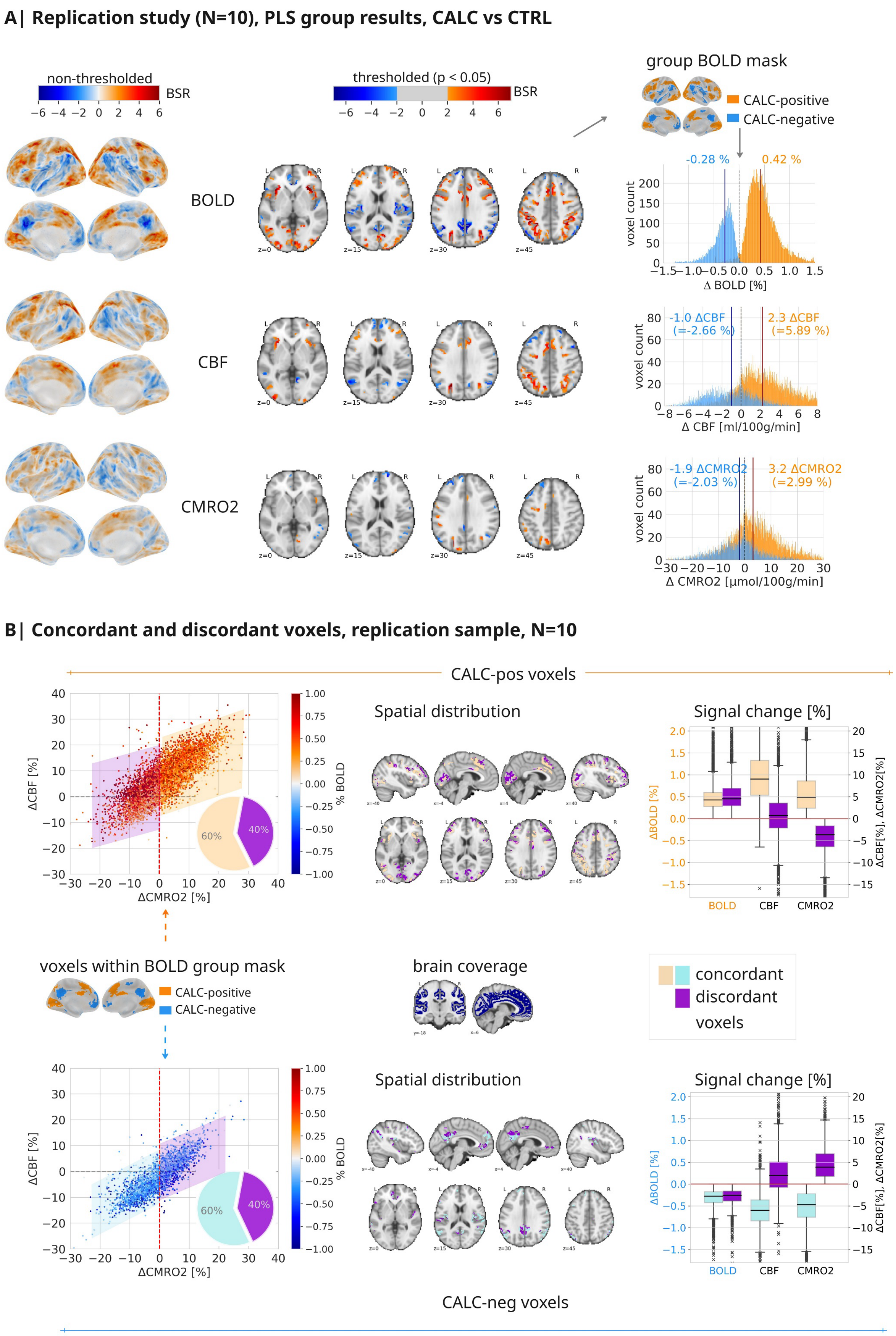


**Figure S6. PLS analyses and concordant versus discordant voxels for a replication sample, N=10. A|** Statistical results of the PLS group analysis in standard space, comparing CALC and CTRL, for both BOLD and quantitative fMRI parameter maps. Maps on the brain surface show non-thresholded statistical maps (values are bootstrap ratios=BSR, akin to z-values) of the first latent variable. The first latent variable was significant for the CBF analysis ($p$ = .03), but not significant for the CMRO2 analysis ($p$ = .05). Red voxels indicate higher values during CALC compared to CTRL, while blue voxels represent the opposite contrast. Axial slices display significant brain clusters, thresholded at BSR ±2 (akin to $p < .05$, cluster size >30 voxels) for BOLD-fMRI, CBF, and CMRO2. Histograms depict across-subject median voxel distributions of ΔBOLD, ΔCBF and ΔCMRO2 (CALC minus CTRL) within the binarized CALC-pos and CALC-neg group masks (shown as surface plot). **B|**CALC vs. CTRL. Data from BOLD and quantitative fMRI for voxels with significant ΔBOLD responses, see group mask on the surface plot. Voxels with ΔBOLD [%] concordant to ΔCMRO2 [%] are depicted in pale (orange and blue), while voxels with discordant ΔBOLD [%] are depicted in violet color. The pie plots summarize the ratio of discordant voxels. Axial slices illustrate the spatial distribution of significant voxels. The boxplots summarize ΔCMRO2[%] and ΔCBF[%] across all voxels with either concordant or discordant ΔBOLD[%]; line, median; box limits, upper and lower quartiles; whiskers, minimum and maximum data points except for outliers: values outside of 1.5* IQR; individual dots, median voxel values). Please note that discordant and concordant voxels feature a similar ΔBOLD amplitude distribution despite indicating opposite metabolic responses. In summary, the replication study shows increased positive and negative ΔCBF and ΔCMRO2 pointing towards a better specificity of the smaller CBF voxels. Nevertheless, the percentage of discordant voxels still amounts to 40% in both CALC-positive and CALC-negative areas, suggesting a substantial mismatch between the sign of the ΔCMRO2 and ΔBOLD responses.

**Replication of Fig. 3 with ΔCMRO2 calculated via the Davis model**

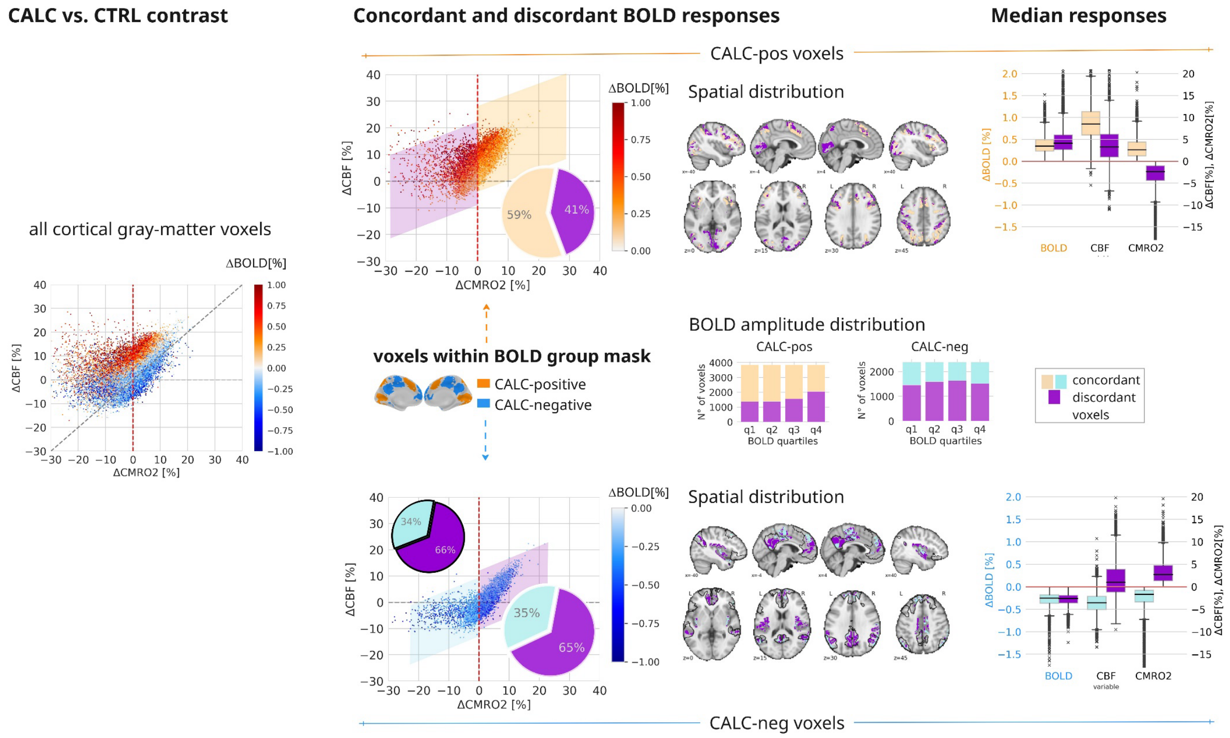


**Figure S7. Replication of Fig. 3 for CALC vs CTRL, with task CMRO2 (CALC) calculated using the Davis model, N=40.** Scatterplots illustrate measured data from both BOLD- and quantitative fMRI for all cortical voxels on the left. In the middle, we separate the significant CALC-negative and CALC-positive voxels. Values are median across all subjects per voxel, colors indicate ΔBOLD [%]. Voxels where ΔBOLD is discordant to ΔCMRO2 are highlighted in violet. The pie charts summarize the ratio of discordant to concordant voxels. Axial slices illustrate the spatial distribution of significant concordant and discordant BOLD voxels. The stacked bars in the right center depict the amplitude distribution of discordant and concordant voxels across BOLD amplitude quartiles. The boxplots summarize ΔCMRO2[%] and ΔCBF[%] across all voxels with either concordant or discordant ΔBOLD[%]; line, median; box limits, upper and lower quartiles; whiskers, minimum and maximum data points except for outliers: values outside of 1.5* IQR; median voxel values across subjects. Please note that discordant and concordant voxels in CALC-positive and CALC-negative masks, respectively, exhibit similar ΔBOLD amplitudes despite reflecting opposite metabolic responses. Compared to Fig. 3 in the main manuscript, the results are very similar: We found 41% (compared of 31%) discordant CALC-positive voxels and 65% (instead of 66%) discordant CALC-negative voxels, alongside comparable spatial distributions and median responses.

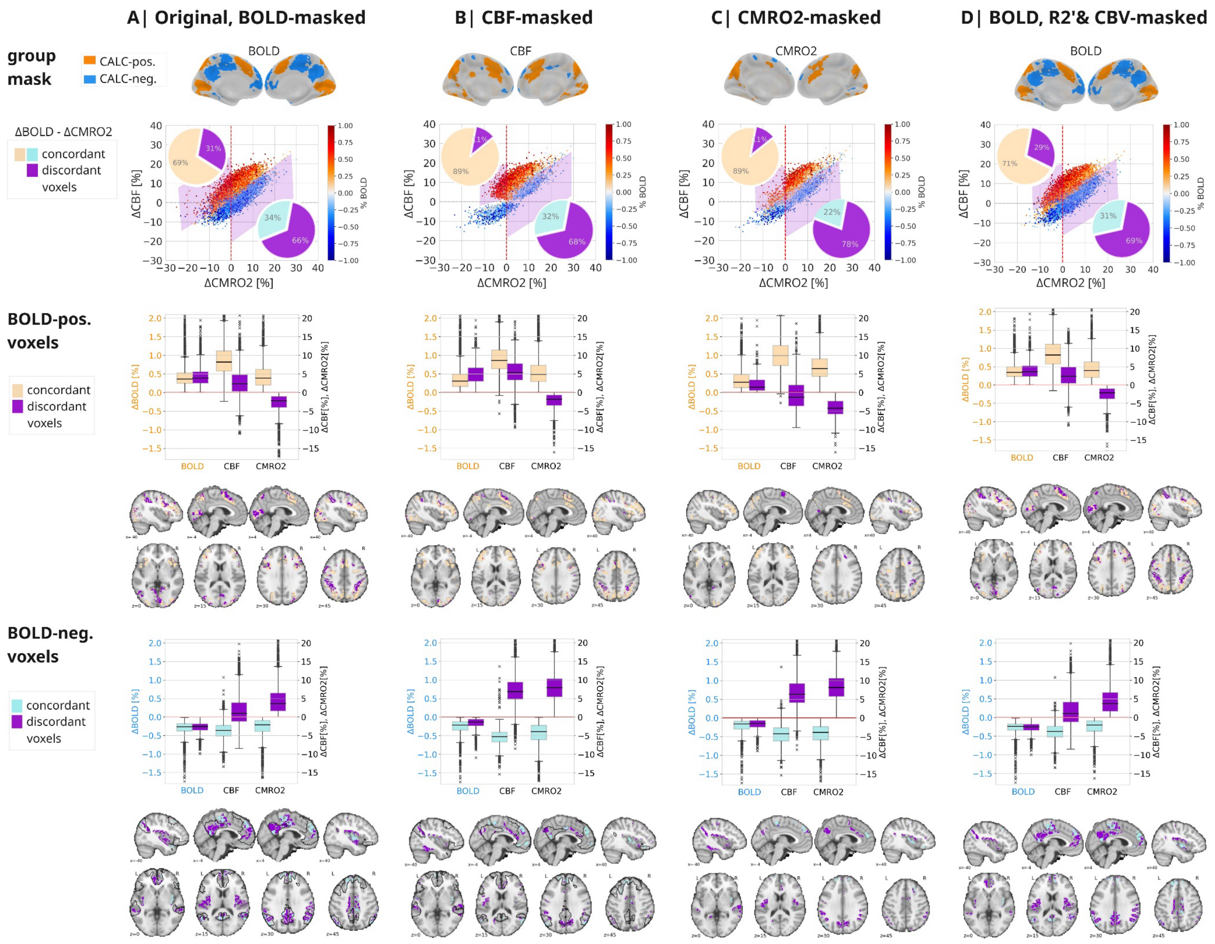


**Figure S8. Dependence of ΔBOLD responses on changes in ΔCBF and ΔCMRO2, CALC vs. CTRL contrast, N=40, with different masks applied.** Scatterplots are showing the subject-averaged ΔCMRO2[%] versus ΔCBF[%] voxels within the respective mask, colors indicate ΔBOLD[%]. Pie plots show the ratio of concordant and discordant voxels separately for positive and negative BOLD voxels. Voxels with ΔBOLD concordant to ΔCMRO2 are depicted in pale (orange = positive ΔBOLD[%], blue = negative ΔBOLD[%]), while voxels with discordant ΔBOLD are depicted in violet color. Axial slices illustrate the spatial distribution of negative and positive BOLD voxels separately, concordant (pale) and discordant (violet) voxels are color-coded. The boxplots summarize ΔBOLD[%], ΔCBF[%] and ΔCMRO2[%] across all voxels with either concordant or discordant ΔBOLD[%], separately for positive and negative BOLD voxels; line, median; box limits, upper and lower quartiles; whiskers, minimum and maximum data points except for outliers: values outside of 1.5* IQR; median voxel values across subjects. **A|** Data as plotted in Fig. 3 in the main manuscript, data were masked by the BOLD group mask. **B|** Same data, yet masked by a group-level CBF mask instead of a BOLD mask, see group CBF mask. While the percentage of discordant BOLD-positive voxels diminished, the percentage of discordant BOLD-negative voxels was still around 2/3 of all BOLD-negative voxels. Please note that in comparison to the original results, ΔCBF[%] across discordant voxels was much stronger, see boxplots. **C|** Same data, yet masked by a group-level CMRO2 mask instead of a BOLD mask, see group CMRO2 mask. While the percentage of discordant BOLD-positive voxels diminished, the percentage of discordant BOLD-negative voxels was even more than 2/3 of all BOLD-negative voxels. Please note that in comparison to the original results, ΔCBF[%] and ΔCMRO2[%] across discordant BOLD-negative voxels was much stronger, see boxplots. **D|** Same data, yet additionally to the group-level BOLD mask, masked by R2’ and CBV, see group mask depicted. All results show only minor changes in comparison to the original results.

**REFERENCES** (Supplementary Information only)


Glasser, M. F., Coalson, T. S., Robinson, E. C., Hacker, C. D., Harwell, J., Yacoub, E., Ugurbil, K., Andersson, J., Beckmann, C. F., Jenkinson, M., Smith, S. M., & Van Essen, D. C. (2016). A multi-modal parcellation of human cerebral cortex. *Nature*, *536*(7615), 171–178. https://doi.org/10.1038/nature18933

Grubb, R. L., Raichle, M. E., Eichling, J. O., & Ter-Pogossian, M. M. (1974). The Effects of Changes in Pa $CO_2$ Cerebral Blood Volume, Blood Flow, and Vascular Mean Transit Time. *Stroke*, *5*(5), 630–639. https://doi.org/10.1161/01.STR.5.5.630

Hyder, F. *et al.* Uniform distributions of glucose oxidation and oxygen extraction in gray matter of normal human brain: No evidence of regional differences of aerobic glycolysis. *J Cereb Blood Flow Metab* **36**, 903–916 (2016). https://doi.org/10.1177/0271678X15625349

Notter, M., Gale, D., Herholz, P., Markello, R., Notter-Bielser, M.-L., & Whitaker, K. (2019). AtlasReader: A Python package to generate coordinate tables, region labels, and informative figures from statistical MRI images. *Journal of Open Source Software*, *4*(34), 1257. https://doi.org/10.21105/joss.01257

Shulman, G. L., FIEZ, J. A., Corbetta, M., Buckner, R. L., Miezin, F. M., Raichle, M. E., & Petersen, S. E. (1997). Common Blood Flow Changes across Visual Tasks: II. Decreases in Cerebral Cortex. *Journal of Cognitive Neuroscience*, *9*(5), 648–663. https://doi.org/10.1162/jocn.1997.9.5.648
